# Skewed temperature dependence affects range and abundance in a warming world

**DOI:** 10.1101/408104

**Authors:** Amy Hurford, Christina A. Cobbold, Péter K. Molnár

## Abstract

Population growth metrics such as *R*_0_ are usually asymmetric functions of temperature, with cold-skewed curves arising when the positive effects of a temperature increase outweigh the negative effects, and warm-skewed curves arising in the opposite case. Classically, cold-skewed curves are interpreted as more beneficial to a species under climate warming, because cold-skewness implies increased population growth over a larger proportion of the species’ fundamental thermal niche than warm-skewness. However, inference based on the shape of the fitness curve alone, and without considering the synergistic effects of net reproduction, density, and dispersal may yield an incomplete understanding of climate change impacts. We formulate a moving-habitat integrodifference equation model to evaluate how fitness curve skewness affects species’ range size and abundance during climate warming. In contrast to classic interpretations, we find that climate warming adversely affects populations with cold-skewed fitness curves, positively affects populations with warm-skewed curves and has relatively little or mixed effects on populations with symmetric curves. Our results highlight the synergistic effects of fitness curve skewness, spatially heterogeneous densities, and dispersal in climate change impact analyses, and that the common approach of mapping changes only in *R*_0_ may be misleading.

## Introduction

Numerous species are undergoing range shifts in response to climate change, typically polewards in latitude or upwards in altitude^1–3^. Underlying these shifts are complex spatial dynamics that may include: regional extirpations in areas where conditions are becoming unsuitable; persistence, but with altered population dynamics in regions where conditions have changed but remain suitable; and dispersal, followed by potential establishment and growth, into regions where conditions have become newly suitable^4,5^. Depending on the respective strengths of these processes, the net effect may be a range shift that corresponds to either an increase or decrease in range size and/or abundance. Examples of such changes exist from almost all major taxa^1–3^, including for some pathogens and pests, as well as for some species that provide ecosystem services^6–9^.

One key element for determining the impacts of a warmer climate on a species’ range and abundance is the temperature sensitivity of its population growth, which in turn is a consequence of the temperature sensitivities of the underlying life history components^10^. Mortality, for example, tends to increase exponentially with temperature within a species’ thermal tolerance range, while fecundity in contrast exhibits a hump-shaped relationship – increasing first to an optimal temperature before decreasing again to zero at high temperatures (see^10^ and references therein). In many insects and vertebrate ectotherms^10–12^, these patterns combine to yield a unimodal temperature-population growth relation that increases gradually to an optimum before dropping steeply to zero near the warmest temperatures of the species’ tolerance range; a relation which we refer to as cold-skewed because the distribution’s heavy tail corresponds to cold temperatures (Figure 1b). By contrast, environmentally transmitted nematode parasites can exhibit a warm-skewed population growth curve, because the mortality of free-living stages is directly affected by temperature, but reproduction – when occurring within an endotherm definitive host – is temperature-independent^13^ (Figures 1c and 2b; see also ref. ^14^). Intermediate cases also exist (e.g. more symmetric population growth curves in bacteria^15^; Figure 1d), and skewness may further vary between^16^ and within taxa^17^. In general, fitness curves are expected to be cold-skewed when the positive effects of a temperature increase (e.g. increased reproduction) outweigh the negative effects (e.g. increased mortality), and are expected to be warm-skewed in the opposite case^18,19^.

**Figure 1.**
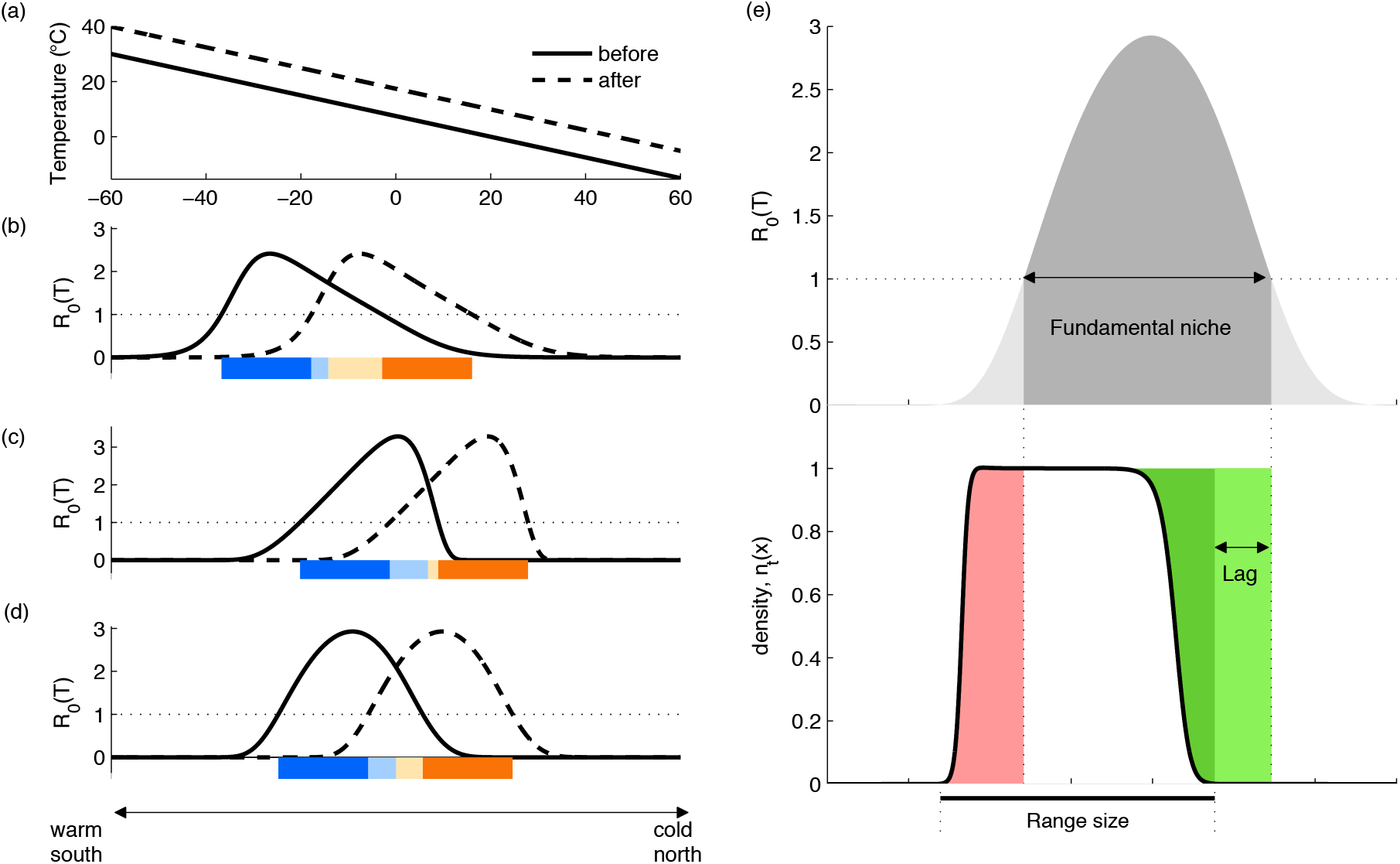
Summary statistics quantifying the impacts of climate warming. (a) The hypothetical south-to-north temperature gradient before (solid line) and after (dashed line) climate warming. (b-d) *R*_0_(*T*) at corresponding spatial locations for cold-skewed (b), warm-skewed (c), and symmetric (d) *R*_0_(*T*). Orange bars indicate increased fitness after warming, with darker orange indicating regions where *R*_0_(*T*)<1 before warming. Blue indicates decreased fitness after warming, with darker blue indicating regions where *R*_0_(*T*)>1 before warming. The percentage of the color bar that is orange (cold-skewed: 57%, warmed-skewed: 44%, symmetric: 50%) is the percentage of the habitat with an increased population growth rate due to climate warming (excluding the regions that are unsuitable both before and after climate warming). (e) illustrates our summary metrics: the ‘fundamental thermal niche’ is defined as all locations where *R*_0_(*T*)>1; ‘range size’ is the length of the spatial domain where the population density is above a numerical cut-off value of 0.001; ‘abundance’ is the numerical integral of the population density across the landscape; ‘lag’ (light green) is the distance between the location of the range front and the front of the fundamental niche; ‘extinction debt’ (the total area of the pink region) is an abundance calculated as the present density integrated across all regions where the population is eventually expected to become extirpated due to *R*_0_(*T*)<1; and ‘colonization credit’ (the total area of the green region) is an abundance calculated as the difference between the present density and the carrying capacity integrated across all regions where the population could persist due to *R*_0_(*T*)>1.

**Figure 2.**
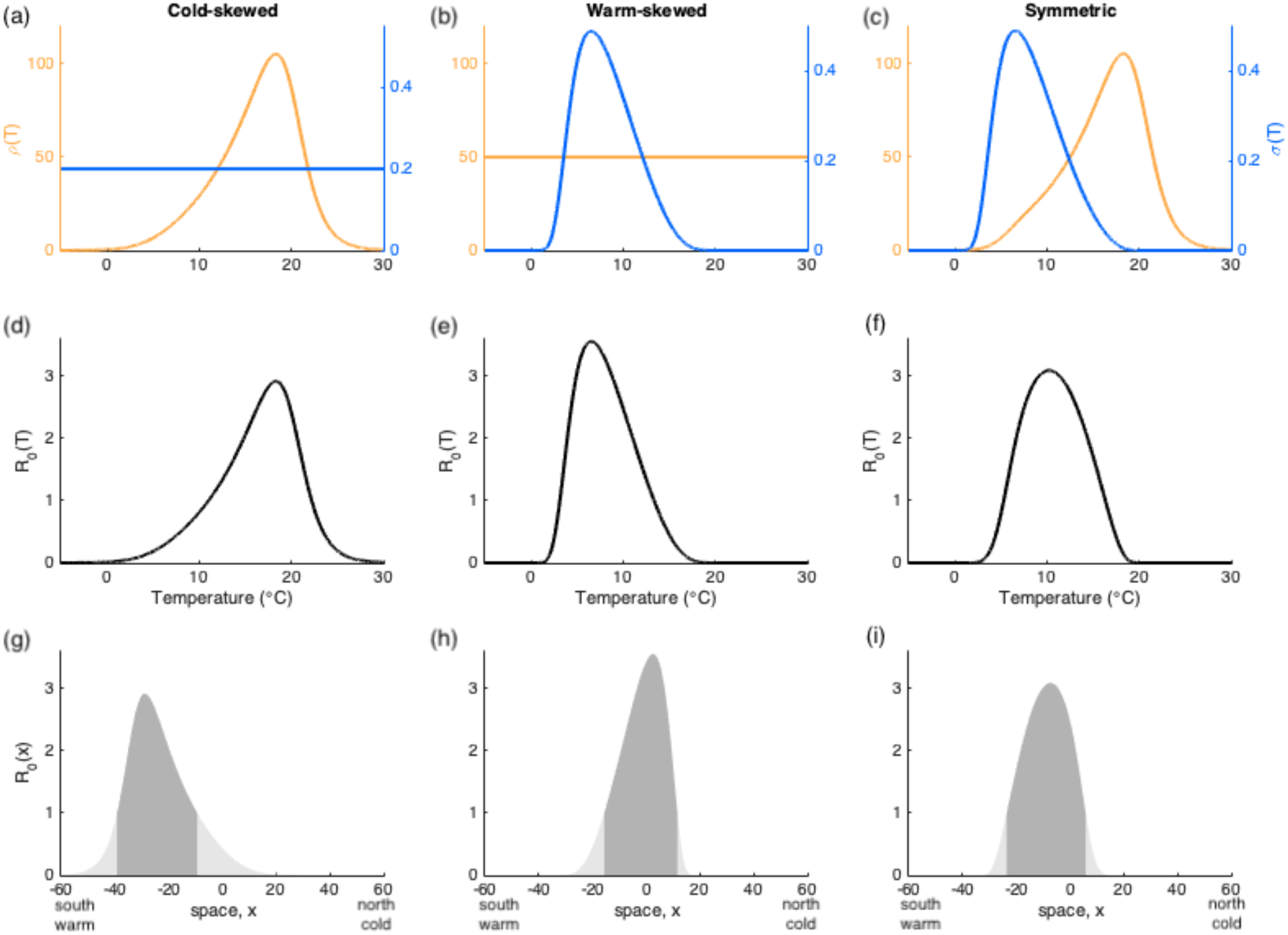
*R*_0_(*T*) curves with different directions of skew arise when fecundity or mortality are more strongly temperature-dependent. (a-c) Sharpe-Schoolfield equations (equations 5–7) describe the temperature dependence of fecundity *(ρ(T),* orange) and survival (*σ*(*T*), blue), yielding (d-f) the temperature-dependent *R*_0_(*T*) as the product of these two quantities (equation 4), which is then normalized so that the sum of *R*_0_(*T*) across the landscape is a constant for niches of all skewness, and (g-i) the spatial dependence of *R*_0_ by combining *R*_0_(*T*) with a spatial temperature gradient (here, linearly decreasing from 30°C at x=-60 in the south to −15°C at x=60 in the north) and with *R*_0_(*T*)≥ 1 shown in dark grey and *R*_0_(*T*)<1 shown in light grey shading. (a,d,g): fecundity only depends on temperature with 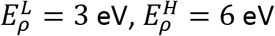, and *μ*(*T*)=*μ*_0_=-ln(0.2); (b,e,h): survival only depends on temperature with 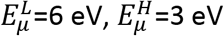, and *ρ*(*T*) = *ρ*_0_ =50 and (c, f, i): both fecundity and survival are temperature-dependent with 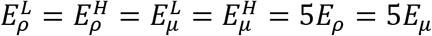. Note that the left and middle columns describe extreme scenarios that generate *R*_0_(*T*)-curves that bound other, more realistic, combinations of temperature dependencies in fecundity and mortality.

The shape of a species’ temperature-dependent population growth curve has been highlighted as a key determinant of where a species is likely to persist in a warming climate. For example, researchers often estimate a species’ fitness curve, e.g., the net reproductive number, *R*_0_, as a function of temperature, and then evaluate whether population growth would increase or decrease in different locations under future climates^11,20–22^, sometimes also comparing the total land area where increases are likely against the total area where decreases are likely as a measure of the anticipated overall net climate change impact^21^. The logical consequence of this perspective is that species with a cold-skewed population growth curve should have an advantage over species with a warm-skewed curve in a warming climate, because a cold-skewed curve maximizes the area over which population growth will increase in response to warming^7,18^ (Figure 1b-d). This interpretation does not, however, explicitly consider the ability of a species to disperse into and colonize new habitats, nor does it account for spatially heterogeneous densities along a species’ range. Here, we show that the explicit consideration of dispersal and population dynamic processes suggests the opposite interpretation, that is, that a warm-skewed population growth curve is more beneficial to a species under climate warming than a cold-skewed curve.

To explore how fitness curve skewness may affect a species’ range size and abundance during climate change-induced range shifts, we formulate a ‘moving-habitat’ integrodifference equation (IDE) model^23^. The IDE framework simultaneously considers both dispersal and net reproduction, and produces ‘travelling wave’ solutions^24^ that describe how a species’ distribution moves across the landscape via reproduction, dispersal, colonization, and mortality. ‘Moving-habitat’ IDE models consider a fundamental niche that shifts due to climate warming, and analyses show that the speed of climate change, the size of a species’ niche, as well as its population growth rate and dispersal ability interact to determine if a species will keep pace with its moving niche^25–27^. Moving-habitat IDE models describe the dynamics of both the shifting niche and the shifting species distribution, and are a suitable framework for understanding when populations will lag behind their fundamental niches as a result of climate warming. Populations that do not disperse will remain in place and ultimately go extinct as their niches shift polewards with climate warming. Moving-habitat IDE models^25–27^, and closely-related partial differential equation models^28,29^, have to date only considered vastly simplified representations of population growth, typically assuming a constant, spatially uniform growth rate within the niche, and spatially uniform declines elsewhere (but see ^30^).

Our model: (i) accounts for temperature dependencies in reproduction and survival using relationships suggested by the Metabolic Theory of Ecology (MTE)^18,31^ to formulate temperature-dependent population growth curves of varying degrees of skewness (Figure 2); (ii) numerically solves the IDE to subject populations to climate change-induced habitat shifts; and (iii) evaluates how skewness affects range sizes, abundances, and the lags between the invasion front and the niche boundaries during climate change (see Figure 1e for definitions). Generally, we find that warming adversely affects populations with cold-skewed fitness curves, positively affects populations with warm-skewed fitness curves, has relatively little or mixed effects on populations with symmetric fitness curves, and that these results are largely robust against different choices of population growth and dispersal mechanisms.

## Methods

IDE models combine reproduction, mortality, and dispersal to describe the spatiotemporal dynamics of a population, treating space and time as continuous and discrete variables, respectively. The population density, *n*_*t*+1_(*x*), at location *x* in year *t*+1 is given by

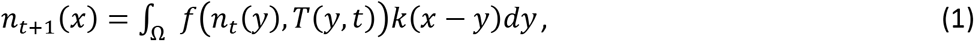

where *f*(*n_t_*(*y*),*T*(*y,t*)) describes the density- and temperature-dependent population growth at location *y* in year *t*, and the dispersal kernel *k*(*x-y*) describes the probability of an offspring dispersing from location *y* to location *x*. We assume that no parents survive after reproduction (i.e., non-overlapping generations). Accordingly, following reproduction, survival and dispersal, the integral totals the number of offspring arriving at location *x* from all possible origins (*y* in Ω) to give the new population density, *n*_*t*+1_(*x*). For simplicity, we model Ω as a one-dimensional domain [*-L,L*], corresponding to either a latitudinal temperature gradient from the equator to a pole, or to an elevation gradient from low to high. For convenience, but without loss of generality, we discuss our model for a latitudinal temperature gradient in the northern hemisphere, hereafter referring to *x*=-*L* as the “south” and *x*=*L* as the “north” and assuming a linear temperature decrease from *y*=-*L* to *y*=*L* (Figure 1a).

In the main text, we focus our discussion on the Beverton-Holt model for population growth, the Laplace dispersal kernel, and deterministic annual temperature increases (Figures 3–4). However, for generality we also evaluate the robustness of our conclusions to other combinations of population growth and dispersal, including compensatory and noncompensatory density dependence and fat-tailed, exponentially bounded, and asymmetric dispersal kernels (ESM Sections 5–7). Moreover, we also consider annual stochastic temperature variations around the deterministically increasing mean (ESM Section 8).

**Figure 3.**
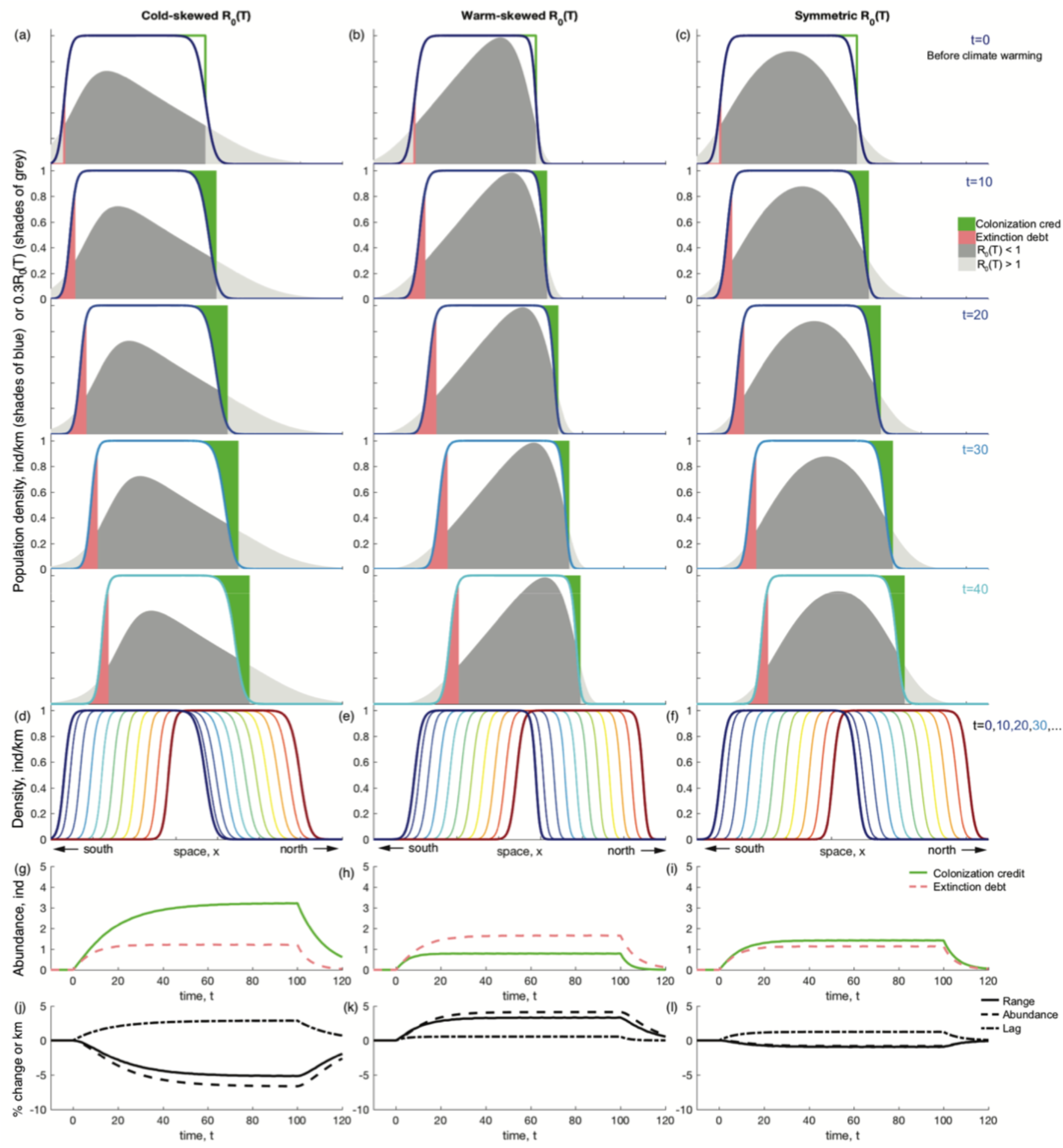
Range change dynamics of species with cold-skewed, warm-skewed and symmetric *R*_0_(*T*) curves. (a-c) At *t*=0, climate warming of +0.1°C yr^−1^ begins causing the *R*_0_(*T*) curve to shift northwards (shaded grey, with regions where *R*_0_(*T*)>1 shown darker). Blue curves show the population density responding to this niche shift. Extinction debt (the abundance of the population persisting in regions where *R*_0_(*T*)<1) is the total area of the shaded pink region and colonization credit (the abundance of the population temporarily below carrying capacity despite *R*_0_(*T*)>1) is the total area of the shaded green region. (d-f) show the population density at 10-year increments during climate warming. Populations with cold-skewed and symmetric *R*_0_(*T*) have colonization credits that exceed extinction debt (pink dashed; g,i) and thus reduced range size and abundance under climate warming (j,l), the opposite is true for populations with a warm-skewed *R*_0_(*T*) (h,k). Parameter values are described in the main text and the ESM Section 1.2, with the activation energies *E_ρ_* and *E_μ_* fixed as *E_ρ_*=0.65 eV and *E_μ_* =0 eV (cold-skewed *R*_0_(*T*)), *E_ρ_* =0 eV and *E_μ_* = 0.65 eV (warm-skewed *R*_0_(*T*)), and *E_ρ_* = *E_μ_* =0.65 eV (symmetric *R*_0_(*T*)), respectively. To allow visual comparison between a population’s *R*_0_(*T*) and its density, (a-c) show *R*_0_(*T*) multiplied by 0.3. Similarly, (j-l) show the lag values multiplied by 50.

The Laplace kernel for dispersal is given by

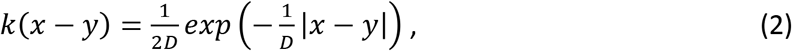

and assumes that the probability of dispersal from location *y* to *x* depends only on the distance |*x-y*| with a mean dispersal distance *D* (see ref. ^32^ for a mechanistic derivation). Specifically, equation (2) assumes that dispersal is not affected by the temperature or density where an individual is born, *y*, where it settles, *x*, or the habitat it travels through to get from *y* to *x*.

The Beverton-Holt model describes annual net reproduction, can be viewed as the discrete time analogue to the continuous time logistic equation, and is given by

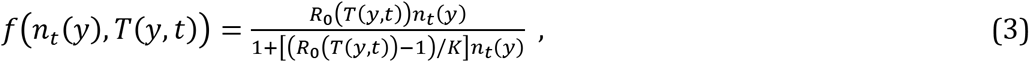

where *K* represents the carrying capacity and *R*_0_(*T*) is the temperature-dependent net reproductive number at low densities, which we refer to as the ‘fitness curve’. Both *R*_0_ and *K* could be impacted by environmental conditions in multiple ways, but we focus only on temperature dependencies in the former to allow disentangling the influence of fitness curve shape on the range change dynamics without confounding these analyses with changes in carrying capacity. We assume a fixed generation time of 1 year (implying that temperature does not affect the number of generations per year) and describe the temperature dependence of population growth via the underlying temperature dependencies of fecundity and mortality,

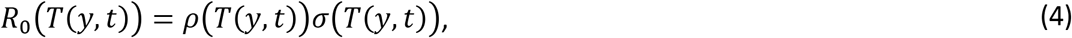

where *ρ* is fecundity at low population density, *σ* is the probability of an individual surviving to reproduce at low population density, and *T*(*y,t*) is temperature at location *y* at time *t*. For notational simplicity, we write *T*(*y,t*)=*T* henceforth.

We describe the temperature sensitivities of both fecundity, *ρ*(*T*), and survival, *σ*(*T*), using relationships suggested by the Metabolic Theory of Ecology (MTE)^18,31^. Fecundity is typically cold-skewed, increasing exponentially with temperature within a species’ fundamental thermal niche, and dropping steeply to zero near the upper and lower temperature boundaries of that range (Figure 2a). Assuming the temperature dependency of fecundity is largely driven by the sensitivity of underlying metabolic processes, we use the Sharpe-Schoolfield model^18^ to represent *ρ*(*T*) and write

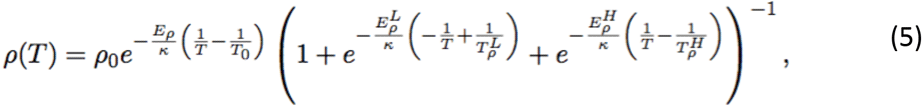

where *ρ*_0_ is the fecundity at a reference temperature *T*_0_ (units: Kelvin, K), κ= 8.62×10^−5^ eV K^−1^ is Boltzmann’s constant (units: electronvolt per Kelvin, eV K^−1^), *E_ρ_* is the average activation energy (units: eV) of the rate-limiting enzyme driving reproduction (determining the temperature sensitivity of fecundity at intermediate temperatures of the organism’s niche), and 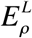 and 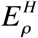 are the inactivation energies (units: eV) determining how abruptly fecundity drops to zero at the thermal tolerance boundaries, 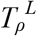 and 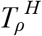.

Survival is typically also a unimodal function of temperature but, unlike fecundity, is usually warm-skewed, peaking near the lower boundary of the organism’s fundamental niche and declining exponentially as temperatures increase (Figure 2b). As with fecundity, we use an

MTE-based formulation^18^, representing the temperature-dependent mortality rate by

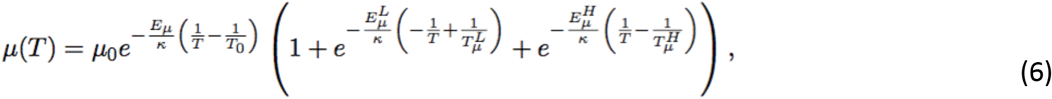

and the proportion of individuals that survive to reproduce by

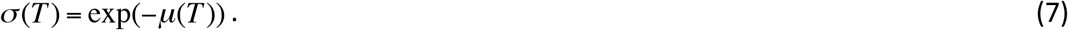

As with fecundity, the parameters 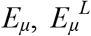 and 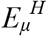 determine the temperature sensitivity of mortality within and outside the lower, 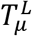, and upper temperature thresholds, 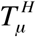, and *T*_0_ is a reference temperature at which mortality equals *μ*_0_.

### Model simulations and parameter values

We assumed a linear temperature gradient ranging from 30°C to −15°C across the landscape (Ω = [–L, L]). To evaluate how the shape of the temperature-dependent fitness curve *R*_0_(*T*) (equation 4) influences the range change dynamics, we establish cold-skewed, warm-skewed, and symmetric *R*_0_(*T*) curves by assuming temperature sensitivities in fecundity only, mortality only, or both (Figure 2, cf. also ref. ^18^). We present simulations for these three extreme cases (Figure 4) and note that they generate a wide variety of *R*_0_(*T*) shapes that bound other, more common, combinations of the relative strengths of the temperature dependencies of fecundity and mortality (i.e. *E_ρ_* ≠ E_μ_ but both positive). We set the inactivation energies (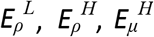, and 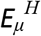) such that they further accentuate the direction of any skew in *R*_0_(*T*) whilst ensuring that the fundamental niche is contained within the spatial domain. The magnitude of the skew is further manipulated in each of these cases by considering a biologically plausible range of activation energies^33^ (0.2 ≤ *E_ρ_, E_μ_* ≤ 1.1 eV). We standardized each fitness curve so that the total reproductive potential of an organism across the entire landscape is always a constant. This calibration ensures that the shape of *R*_0_(*T*) is the main source of variation in all comparisons, and that qualitative differences in climate change impacts can be attributed to differences in the fitness curve shape (ESM, Section 2). We used numerical simulations to explore the range change dynamics given by equations 1–7. Population density was allowed to equilibrate before the onset of warming (at t=0), after which we increased the temperature at all locations by *w*=0.1°C yr^−1^ until an increase of 10°C was achieved. We tracked the population’s range size and abundance, as well as the lag between the northern boundary of the population range and of its thermal niche (see Figure 1e for definitions). Range size and abundance changes are reported as percentage changes relative to the range and abundance at the onset of warming (*t*=0), whereas lag is reported as the difference between future lags and the lag at *t*=0 to facilitate comparisons between fitness curves with different shapes. The MATLAB code is provided on Figshare^34^ (doi: 10.6084/m9.figshare.6955370) and further details of the simulation parameterization, assumptions, and implementation are described in the ESM (Sections 1 and 2).

**Figure 4.**
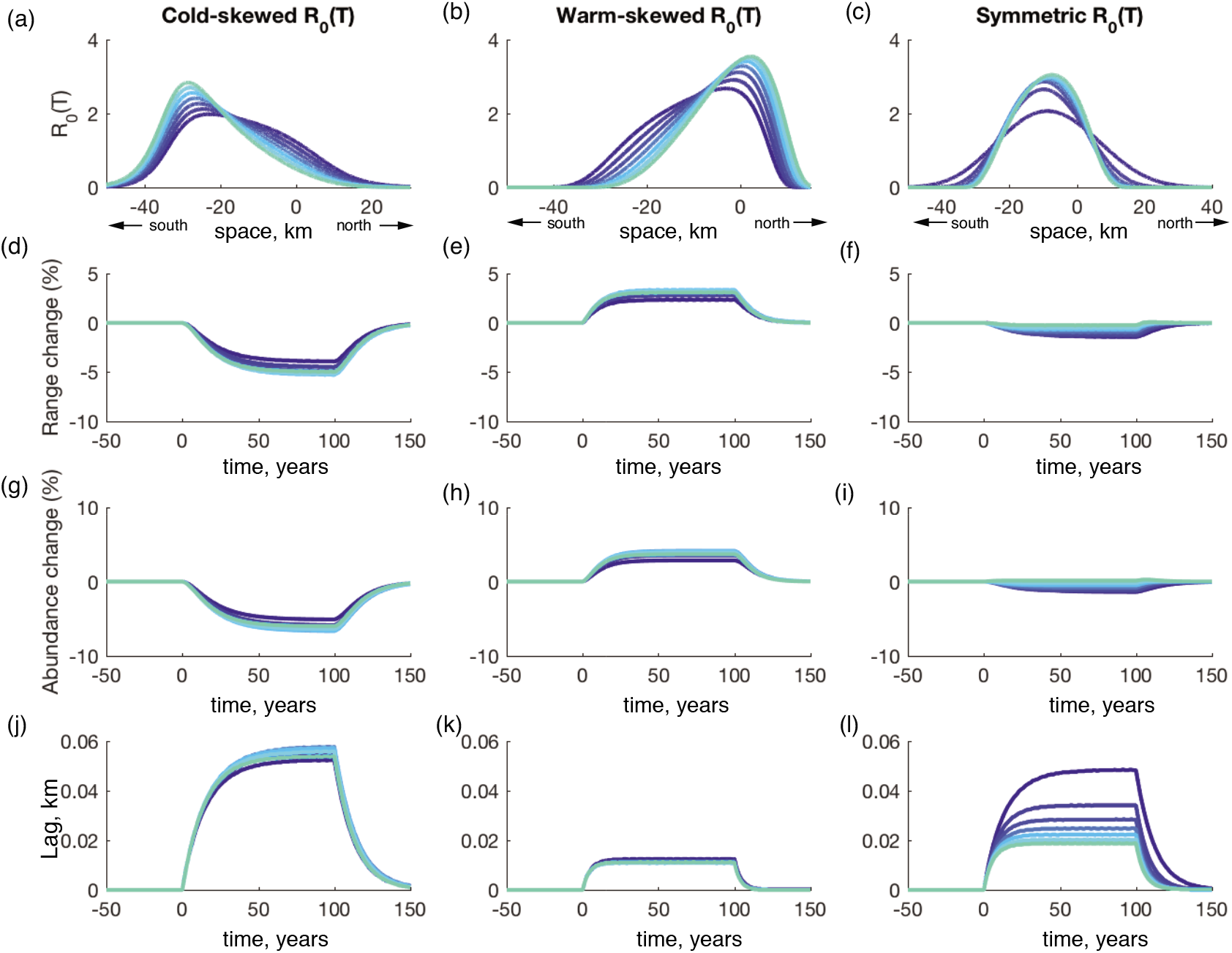
Climate warming adversely affects populations with a cold-skewed *R*_0_(*T*) (left column), benefits populations with a warmed-skewed *R*_0_(*T*) (middle column), and has relatively little effect on populations with a symmetric *R*_0_(*T*) (right column). (a-c) Different degrees of *R*_0_(*T*) skewness are explored for each of the cold-skewed, warm-skewed, and symmetric scenarios by varying *E_ρ_* and/or *E_μ_* from 0.2 to 1.1 eV by steps of 0.15 eV (dark to light blue) (all other parameters are given in the main text and the ESM Section 1.2). (d-l) Changes in range size, abundance, and lag before (*t*_0_), during (*t*=0 to *t*=100), and after (*t*>100) warming. The vertical axes in (d-i) are percentage changes relative to the initial range and abundance and in (j-l) are the absolute difference between future lags (*t*>0) and the lag at *t*=0 (cf. ESM Section 1 and 2 for details).

## Results

Solving equations 1–7 with simulated climate warming reveals that the species’ distribution lags behind its shifting fundamental niche (see also ^5^). We observe regions of ‘colonization credit’^4^, where suitable habitat (*R*_0_(*T*)≥1) has yet to be fully colonized, and of ‘extinction debt’^4,35^, where individuals continue to persist temporarily in unsuitable habitat (*R*_0_(*T*)<1; Figure 3a-c; see Figure 1e for our definitions of colonization credit and extinction debt).

During climate warming, population density retains its general shape, but the skew of *R*_0_(*T*) affects the length of habitat where the population is at carrying capacity (Figure 3a-f). Populations with a warm-skewed *R*_0_(*T*) have small lags (Figures 3k and 4k) and small colonization credits (Figure 3h), which means that most of the fundamental niche has been fully colonized, and suggests increased range sizes (Figures 3k and 4e) and increased abundances (Figures 3k, 4h) under climate warming. The opposite holds for cold-skewed *R*_0_(*T*), where a larger lag (Figures 3j, 4j) results in large colonization credits (Figure 3g), meaning that a large region of the fundamental niche has yet to be fully colonized, and leads to decreased range sizes (Figures 3j, 4d) and abundances (Figures 3j, 4g).

These differences in the range change dynamics of warm-skewed, symmetric, and cold-skewed *R*_0_(*T*) can be understood by considering the population’s potential for range expansions at the leading range edge. Populations with warm-skewed *R*_0_(*T*) are sensitive to beneficial temperature increases in the north, where the slope of the *R*_0_(*T*) curve is steep. In addition, warm-skewed populations have higher densities at the leading edge of their fundamental niche (Figure 3b) due to smaller lags (Figure 4k), and thus a large potential for colonizing newly available northern habitats via dispersal, resulting in an increased range size and abundance during warming (Figure 4e,h). For populations with a cold-skewed *R*_0_(*T*), these same mechanisms act in the opposite direction, implying decreased range sizes and abundance. Extinctions at the trailing edge of the population density also determine the net impact of climate warming, with extinction debts largest for populations with warm-skewed *R*_0_(*T*), further explaining why these populations show increased range size and abundance in response to climate warming (Figures 3g-i; ESM Sections 4 and 5.1). Populations with a symmetric *R*_0_(*T*) are mostly unaffected by climate warming, but show slight decreases in both range size and abundance (Figure 4f,i). These slight decreases occur where the population lags behind its fundamental niche, resulting in a negative net effect of colonization versus extinction, despite the symmetry of *R*_0_(*T*).

The influence of *R*_0_(*T*)’s skewness on a population’s response to climate warming is largely insensitive to aspects of the fitness curve shape other than skewness, mean dispersal distance, the choice of population growth function, the choice for dispersal kernel, and annual stochastic fluctuations in temperature (ESM Sections 3–8; summarized in Table 1), however, for large values of *R*_0_(*T*), Ricker growth may give rise to oscillatory or chaotic travelling waves that do not occur with Beverton-Holt growth^26^, and these oscillations can be dampened by climate warming due to the temperature dependence of *R*_0_(*T*) (ESM Section 6).

**Table 1.** The skewness of *R*_0_(*T*) has a strong influence on a population’s response to climate warming. These effects are largely insensitive to choice of population growth function, dispersal kernel, mean dispersal distance, and climate stochasticity. In response to 100 years of warming, range size and abundance generally decrease (-) with a cold-skewed *R*_0_(*T*), increase (+) with a warm-skewed *R*_0_(*T*), and remain relatively unchanged (0) with a symmetric *R*_0_(*T*).

| Scenario | Population metric | Response after 100 years of climate warming |  |  | Additional information |
| --- | --- | --- | --- | --- | --- |
|  |  | Cold skew | Warm skew | Symm. |  |
| <u>Case 1</u><br>• Beverton-Holt<br>• Laplace | Range size | - | + | 0 | Results are insensitive to variation in mean dispersal distance, $D$ (equation 2; Figures S6 and S7) and to considering asymmetric instead of symmetric dispersal (Figure S8). |
|  | Abundance | - | + | 0 |  |
| <u>Case 2</u><br>• Ricker<br>• Laplace | Range size | - | + | 0 | Results are indistinguishable from Case 1 for small values of $R_0(T)$ (Figures S9 and S10). For large values of $R_0(T)$, population cycles may appear and climate change influences the amplitude of these cycles (Figure S11). |
|  | Abundance | - | + | 0 |  |
| <u>Case 3</u><br>• Beverton-Holt<br>• Cauchy | Range size | 0 | + | + | Figures S10 and S12. |
|  | Abundance | - | + | - |  |
| <u>Case 4</u><br>• Ricker<br>• Cauchy | Range size | 0 | + | + | Figure S10. |
|  | Abundance | - | + | 0 |  |
| <u>Case 5</u><br>• Stochastic fluctuations in annual temperature | Range size | - | + | - | Figure S13. |
|  | Abundance | - | + | - |  |

## Discussion

The shape of temperature-dependent population growth measures such as *R*_0_(*T*) contains critical information on a species’ sensitivity to climate change. The common approach of only considering how population growth metrics would change in different locations under climate change (e.g. *R*_0_(*T*)-or *r*(T)-based impact maps for insects^11^, wildlife pathogens^22^, and human pathogens^20,21^; where *r*(T) is the net reproductive rate at low densities), however, can yield an incomplete and potentially misleading picture. In contrast to the suggestion by Molnár *et al.* (2013)^18^ that cold-skewed populations experience increased population growth over the majority of their habitat (orange bars in Figure 1b) and warm-skewed populations experience increased growth over only a small portion of their habitat (orange bars in Figure 1c), when embedding *R*_0_(*T*) within a framework that considers dispersal, density dependence and a shifting fundamental niche, we find that generally climate warming adversely affects the range size, abundance and lag of populations with a cold-skewed *R*_0_(*T*), positively affects populations with a warm-skewed *R*_0_(*T*), and has relatively little or mixed effects on populations with a symmetric *R*_0_(*T*) (Figure 4; Table 1). For cold- and warm-skewed *R*_0_(*T*), the degree of the skew further magnifies these effects (Figure 4). While previous interpretations of the effect of fitness curve skewness on species’ responses to climate warming have considered solely the shape of *R*_0_(*T*), we suggest that it is not only the skewness of *R*_0_(*T*), but also the synergistic effects of dispersal and spatially varying population densities that underly our findings that a warm-skewed *R*_0_(*T*) results in range size and abundance increases, and small lags between the moving cold tolerance limit of the fundamental niche and the population’s leading edge, whereas a cold-skewed *R*_0_(*T*) results in range contractions, decreased abundances, and large lags.

Our model is deliberately simple to facilitate a sound understanding of the synergistic effects of fitness curve skewness, density, and dispersal as a previously unrecognized mechanism for why some species may benefit from warming while others experience range contractions and population declines. These synergistic mechanisms should be viewed as complementary to previously recognized influences, such as the role of interspecific interactions^37^ in confining species to only part of their fundamental niche. The muskox lungworm *Umingmakstrongyls pallikuukensis*, for example, has a warm-skewed *R*_0_(*T*)^13^ and has been closely tracking the northern boundary of its shifting fundamental niche during a warming-induced range expansion^6^ as would be expected (Figures 3–4). No changes were reported in the southern boundary of the lungworm’s distribution^6^, which is consistent with our results that warm-skewed *R*_0_(*T*) have large extinction debts (Figure 3h), and may further be explained by the behavioral thermoregulation of the worms’ intermediate slug host and the shelter it provides from lethal high-temperature extremes through behavioural thermoregulation^13^. Also in agreement with our results are the range contractions observed in tree species in the eastern United States^38^ and southern Africa^39^, where limited dispersal has resulted in large dispersal lags at the poleward limits, perhaps further exacerbated by apparently cold-skewed fitness curves (cf. Figure 3 in Talluto et al.^4^). Lepidopterans^8,9^ and Odonata^40^, by contrast, have largely expanded their ranges polewards in response to warming – often while maintaining stable or more slowly moving equatorward boundaries – despite generally cold-skewed fitness curves (see Deutsch et al. 2008^11^, their Dataset S1). It has, however, been suggested that these range change patterns could be a result of temperature limiting species’ distribution at the poleward boundary, and factors other than temperature (e.g. competition) currently limiting distribution at the equatorward boundary^8,9,40^. In other words, these species may not be fully occupying the warmest end of their fundamental thermal niche, and would temporarily be shielded from the worst consequences of warming despite having a cold-skewed fitness curve.

The dynamics of climate warming-induced range changes are complex and our analyses are not meant to imply that skewness will uniquely determine whether warming benefits or harms a species. Dedicated meta-analyses are needed to disentangle the various factors affecting range changes (i.e., sensitivities to other abiotic variables^41^, biotic interactions^42^, and landscape features^43,44^), but are currently limited by: (i) unknown fitness curve shapes for most species; (ii) the difficulty of inferring fitness curve shape from local densities when carrying capacity is temperature-independent as assumed here (cf. Figure 3a-f, showing ‘rectangular travelling waves with soft edges’ regardless of fitness curve skew); and (iii) a paucity of studies^45^ documenting distribution and abundance changes over a species’ entire range and on long-enough time scales to capture the dynamics suggested here. Nevertheless, given the robustness of our results to different forms of density dependence and dispersal (Table 1), we expect that imbalances in the temperature sensitivities of different life history traits, and the resultant fitness curve skewness, may explain some of the observed variation in distribution and abundance change responses to climate warming.

Our models can be easily extended in multiple ways by relaxing simplifying assumptions or adding additional population regulating mechanisms. The list of potential factors influencing range change dynamics is long, and includes potential temperature dependencies in other life history and population dynamics parameters (e.g. increased temperature could imply: decreased development times due to faster metabolism^31^; shortened dispersal distances due to decreased larval development times^31^ (e.g. in marine plankton^46^); and decreased carrying capacity, since faster metabolism accelerates the depletion of a fixed resource supply^12^), local adaptation (e.g. different temperature sensitivities of individuals at the leading and trailing edge^17,47^), limiting abiotic factors other than temperature^41^ (e.g. moisture), age-structure^26^ (e.g. for species where dispersal only occurs in certain life stages such as trees or many insects), Allee effects limiting range shift speeds at the leading edge^24,48^, as well as landscape heterogeneities^43,44^, and species interactions^28,37,42^. Each of these might dampen or amplify the patterns suggested by our analyses.

We used temperature-dependent relationships for fecundity and survival but did not explore the effects of temperature dependence on generation time. This assumption most closely corresponds to organisms with a fixed, one-year generation, for example driven by obligatory diapause to overwinter a cold season (e.g. Lepidoptera in temperate forests). However, even in these cases, in response to warmer temperatures, generation times may decrease to allow for multiple generations within a year. Such changes would not affect *R*_0_(*T*) (because *R*_0_(*T*) measures population growth per generation)^10,49^, but would shift the relationship between temperature and the geometric growth rate, *λ*(*T*), towards more cold-skewed, likely with the corresponding consequences for abundance and range changes that were outlined in our Results section. Our combined MTE-IDE framework allows easy incorporation of such additional factors, but would need to be extended to consider multiple generations per year (i.e., ref. ^50^).

The MTE describes temperature effects from the individual to the ecosystem level^31^ and has provided many insights describing how a warmer climate may alter population and community dynamics^18,33,44,51^, but analyses have generally not explicitly considered dispersal. Likewise, there exists a rich literature that uses integrodifference equation models to explore how various intrinsic and extrinsic mechanisms affect range change dynamics^24,26,32,43,44,48^, but few studies have explicitly explored the role of temperature in these dynamics. Combining these two bodies of theory has enormous potential for unravelling the complexities of temperature-dependent spatial population dynamics.

## Acknowledgements

This research was catalyzed by a Banff International Research Station workshop (13w5095) and further facilitated by an Atlantic Association for Research in the Mathematical Sciences Collaborative Research Grant. AH (RGPIN 2014-05413) and PKM (RGPIN 2016-06301) were supported by National Sciences and Engineering Research Council of Canada Discovery grants, and PKM was further supported by the Canada Foundation for Innovation John R. Evans Leader Funds, and MRIS Ontario Research Funds. CC was supported by the Biotechnology and Biological Sciences Research Council (BB/P004202/1). Part of this work was carried out while CC was a funded visiting scholar at the Fields Institute for Research in Mathematical Sciences.

## Supplementary Material

### 1 Additional simulation details and parameter values

The main text provides a general overview describing the simulation of our moving-habitat inte-grodifference equation model with spatially non-uniform net reproduction (equations 1–7). Additional specific details are provided in this section and the MATLAB code used to generate all simulations is archived at Figshare (Hurford et al. 2018; doi: 10.6084/m9.figshare.6955370).

#### 1.1 Simulation details

Temperature is modelled as a linear gradient along the spatial domain [–*-L, L*] that peaks at *y* = -*L*, and warming is simulated by increasing temperatures at all locations at a constant rate:

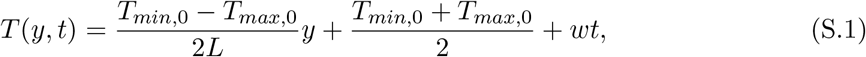

where *T*_*max*,0_ and *T*_*min*,0_ are the respective temperatures at the southern (*y* = –*L*) and northern (*y* = *L*) limits of the spatial domain at time *t* = 0, and *w* is the rate of temperature increase. For simplicity, we ignored seasonal temperature fluctuations, assuming a constant temperature for any fixed location within each year (Figure 1a).

We used computer simulations to explore the range change dynamics given by equations 1–7 in the main text. To do this, we set the initial abundance to 1 on a small interval about *y* = 0, and calculate population densities across the entire spatial domain for 400 years with no change in climate (*w* = 0) to let the population density equilibrate before the onset of warming. Subsequently, we increase the temperature at all locations by *w* = 0.1°C yr^−1^ up until an increase of 10°C is achieved after 100 years, and evaluate how warming alters a population’s range and abundance, as well as the lag between the northern boundary of the population range and the northern boundary of its thermal niche (see Figure 1 in the main text for the definition of the summary metrics). We recognize that a warming of 10°C is far more than expected under current climate change predictions, but note that this value was simply chosen to allow easy visualization of how climate change impacts range, abundance, and lag. All results are qualitatively insensitive to the total amount of warming imposed.

To evaluate how the shape of the temperature-dependent fitness curve *R*_0_(*T*) (equation 4) influences the range change dynamics, we establish cold-skewed, warm-skewed, and symmetric *R*_0_(*T*) curves by assuming temperature sensitivities in fecundity only, mortality only, or both (cf. also ref. Molnar et al. 2013). The strongest cold-skew in *R*_0_(*T*) arises in populations with highly temperature-dependent fecundity (large *E_ρ_*), but temperature-independent mortality (*E_μ_* = 0 eV). A strongly temperature-dependent mortality (large E_μ_) combined with temperature-independent fecundity (*E_ρ_* = 0 eV), by contrast, leads to a strongly warm-skewed *R*_0_(*T*), and equal temperature sensitivities (*E_ρ_* = *E_μ_*) yield an approximately symmetric *R*_0_(*T*) (Figure 2). We present simulations for these three extreme cases and note that they generate a wide variety of *R*_0_(*T*) shapes that bound other, more common, combinations of *E_ρ_* and *E_μ_* (i.e. where *E_ρ_* ≠ *E_μ_* but both > 0 eV; Figure 4).

#### 1.2 Parameter values

The three cases of cold-skewed, warm-skewed, and symmetric *R*_0_(*T*) are implemented by considering temperature dependence in (i) fecundity only (*E_ρ_* > 0, *E_μ_* = 0 eV), (ii) survival only (*E_ρ_* = 0, *E_μ_* > 0 eV), and (iii) both in fecundity and survival (*E_ρ_* = *E_μ_* > 0 eV). The magnitude of the skew is further manipulated in each of these cases by considering a biologically plausible range of activation energies (Dell et al. 2011; 0.2 ≤ *E_ρ_, E_μ_* ≤ 1.1 eV), and by setting the inactivation energies such that they further accentuate the direction of any skew in *R*_0_(*T*) whilst ensuring that the fundamental niche is contained within the spatial domain (case (i): 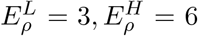 eV; case (ii): 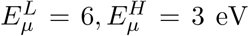 eV; case (iii): 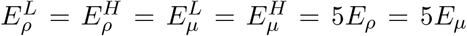; Figure 4). The mean dispersal distance is required to be large enough so that the population can keep pace with its moving niche and was set as *D* =0.5 km. The remaining model parameters were fixed arbitrarily, but the dispersal parameter is required to be large enough so that the population can keep pace with its moving niche (*K* = 1 individuals km^−1^, *μ*_0_=-ln(0.2), *p*_0_ = 50, 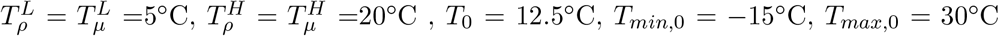; note that all temperatures are reported in °C for simplicity but converted to Kelvin for use in equations 5–6). Finally, we standardized each fitness curve so that the total reproductive potential of an organism across the entire landscape is always a constant (chosen as 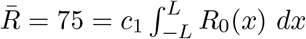 where *c*_1_ is the calibration parameter). This calibration ensures that the shape of *R*_0_(*T*) is the main source of variation in all comparisons, and that qualitative differences in climate change impacts can therefore be attributed to differences in fitness curve shape (see Section S2). While the values of the reference temperature for reproduction and survival, *T*_0_, the reproduction, *ρ*_0_, and mortality rates, *μ*_0_, at the reference temperature, the tolerance limits for reproduction, 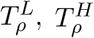, and survival, 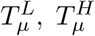, as well as the calibration parameter, *c*_1_, were chosen arbitrarily, our results are not sensitive to these choices because it is the shape of the fitness curve, rather than the specific details of how the curves are produced, that determines the impact of climate warming.

### 2 Scenario standardization

#### 2.1 How we standardize comparisons between fitness curves with different skewness

Throughout this study, we quantify the impacts of climate warming by comparing range size, abundance, and lag relative to their initial values. The range change (%) is calculated by taking the range size at time *t* divided by the initial range size (when climate warming begins at *t* = 0) and multiplied by 100, and the abundance change (%) is calculated in the same way. The lag is calculated by subtracting the lag at time *t* = 0 from the lag at time *t*. Final range size, final abundance, and final lag are calculated as described above, but with *t* = 100 (the time when climate warming ends). In addition, we normalized all fitness curves so that the reproductive potential of a population (the integral of *R*_0_(*T*) across all of space) is equal to the same value 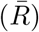 regardless of fitness curve skew. This normalization was necessary to ensure that results reflect differences in fitness curve skewness and are not due to associated changes in reproductive potential.

#### 2.2 Differences in the initial range size and initial abundance cannot explain the observed impact of climate change

The normalization of the reproductive potential for all fitness curves to 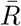 does not ensure that the initial range sizes, initial abundances, or initial lags are also equal for the different choices of *R*_0_(*T*) (Table S.1). In Figure S.1, we demonstrate that differences in initial range size and initial abundance between different *R*_0_(*T*) curves cannot explain the climate induced changes in the population metrics, instead these are driven by the differences in fitness curve skewness. To interpret Figure S.1, note that fitness curves with cold-skew (*), warm-skew (Δ), and symmetric (•) shapes may all have initial range sizes close to 41 (left column), but the initial range size of 41 corresponds to a wide range of final range sizes (between –6 and +2% change), whereas cold-skewed curves (*), of any initial range size, correspond to a narrower range of final range sizes (between +2 and +4% change; Figure S.1a). This pattern is generally consistent for different initial abundances (right column), and if different climate warming impacts are considered (rows), and so we conclude that differences in initial range size or initial abundance are not responsible for the different climate warming impacts that we observe.

**Table S.1:**
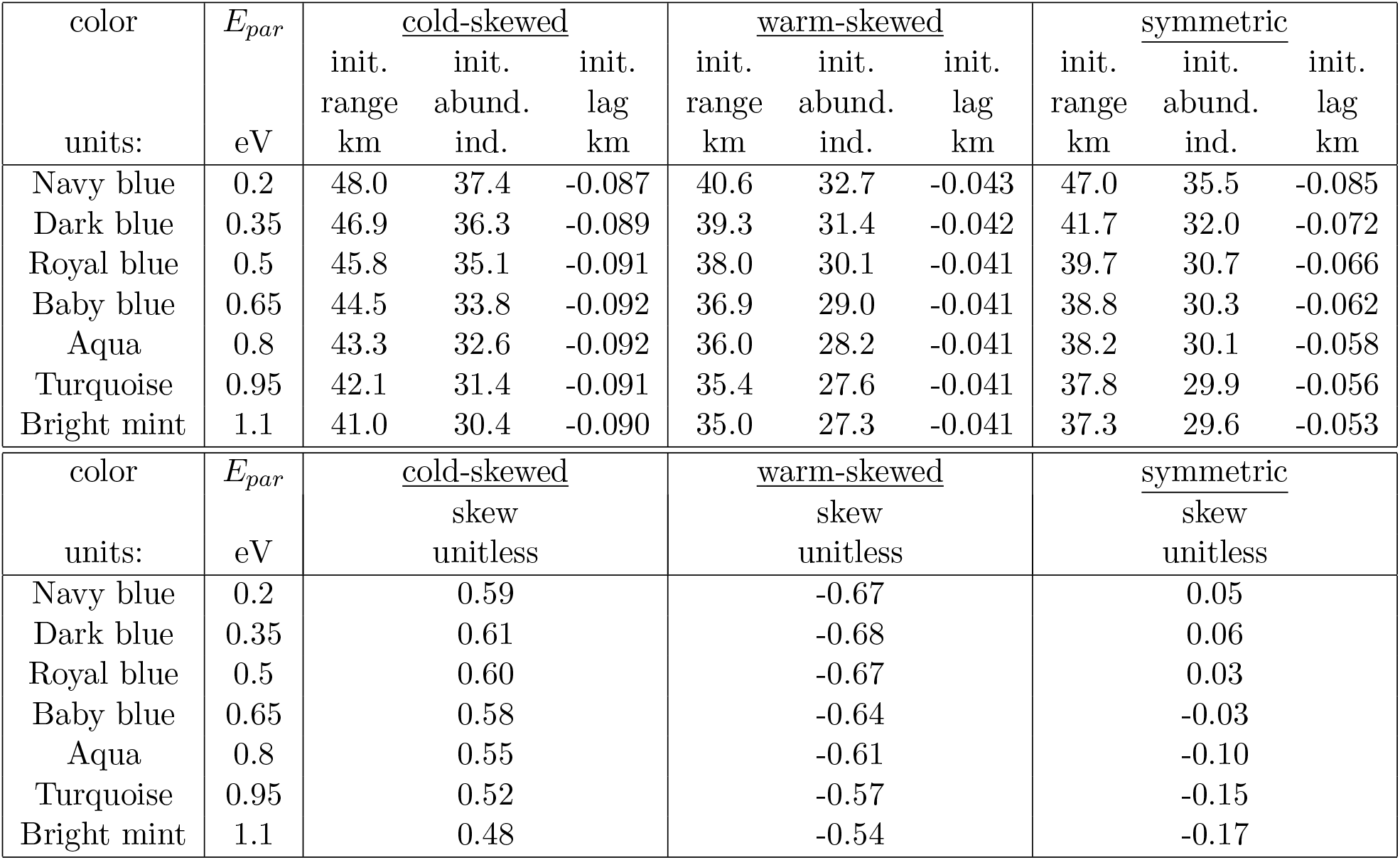
The initial range size, initial abundance, initial lag and skew for all the fitness curves considered in Figure 4 of the main text. All fitness curves are normalized so that the reproductive potential (the integral of *R*_0_(*T*) across all of space) is the same regardless of fitness curve skew, but this does not lead to equal initial range sizes, initial abundances and initial lags. *E_par_* is the parameter that is manipulated to adjust the strength of the skew: *E_par_* = *E_ρ_* (cold-skewed), *E_par_* = *E_μ_* (warm-skewed), and *E_par_* = *E_ρ_* = *E_μ_* (symmetric). Negative values of the lag mean that the population density is above the detection threshold and the population is persisting in a region ahead of the thermal tolerance limits described by the fundamental niche. Negative lags are due to a continuous space analog of the familiar source-sink dynamics described for discrete space patch-based metapopulation models (Pulliam, 1988), whereby a population outside the fundamental niche is sustained through dispersal. Skew is calculated by converting the fitness curves to probability densities (i.e., dividing *R*_0_(*T*) by the integral of *R*_0_(*T*)) and using the equation for Pearson’s first skewness coefficient calculated with respect to space, *x*. A positive skew indicates that the tail of the distribution is in the positive x direction, corresponding to cold temperatures. As such, positive values indicate cold-skewed fitness curves.

**Figure S.1:**
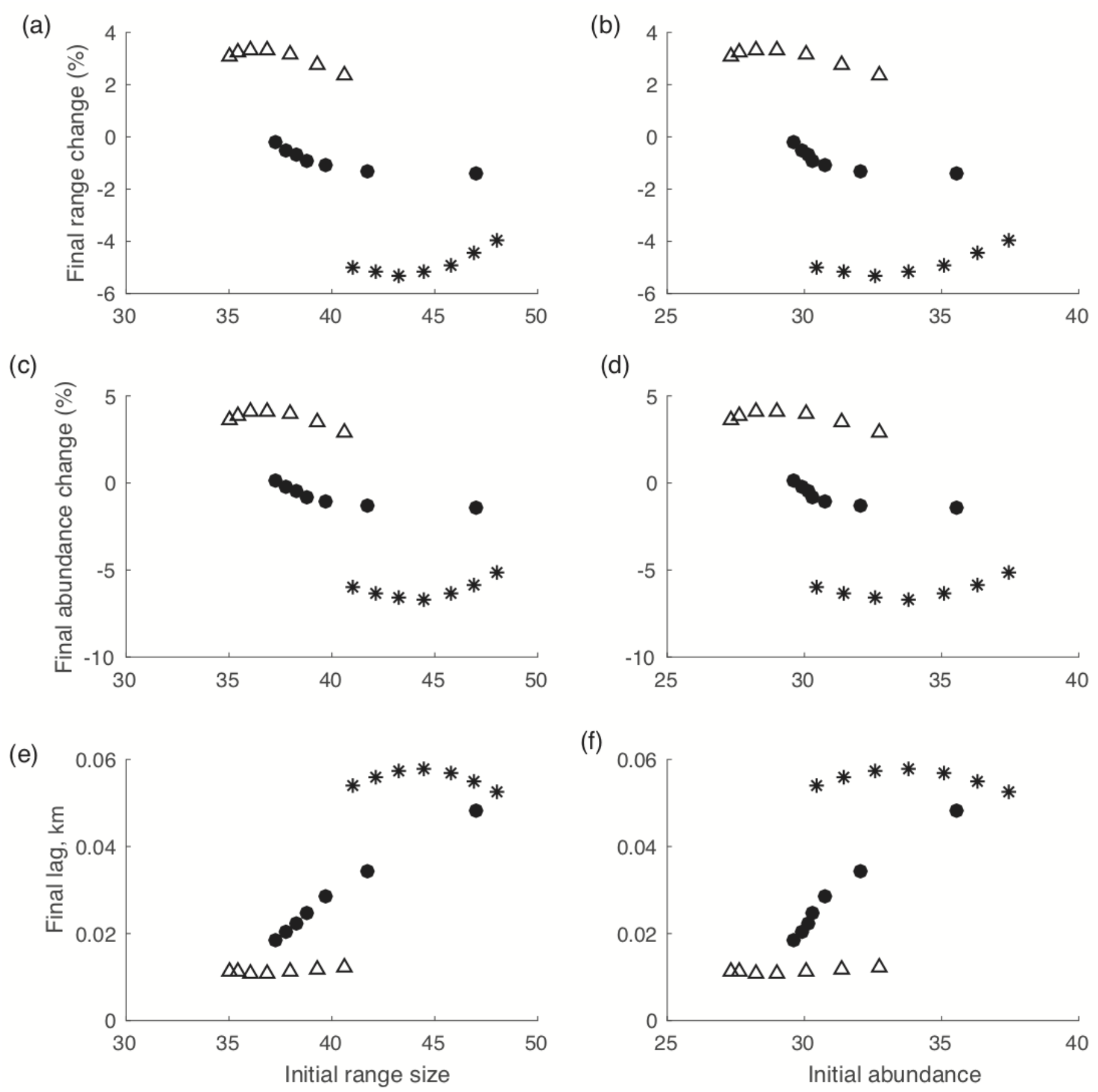
Initial range size (left column) and initial abundance (right column) do not have a strong effect on how warming impacts final range size (top row), final abundance (middle row) and final lag (bottom row). In contrast, there is a strong relationship between the skewness of *R*_0_(*T*) and the effects of climate warming (symbols: cold-skewed – *, warm-skewed – Δ, symmetric – •).

### 3 Fitness curve shape

#### 3.1 Kurtosis

The kurtosis of the fitness curves we consider is imposed by equations 5–6 and their parameterization, particularly the inactivation energy parameters (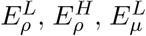 and 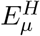), which determine how abruptly the fitness curves decrease beyond their thermal tolerance limits. Figure S.2 (right column) illustrates the kurtosis of each fitness curve considered in Figure 4 of the main text (top row) plotted against the effects of climate warming. The effect of kurtosis is much less important than the effect of skewness. The correlation coefficients for skewness relative to range size (Figure S.2a) and abundance change (Figure S.2c) are both *R*^2^ = −0.99, whereas the correlation coefficients for kurtosis relative to these same quantities are *R*^2^ = −0.67 and −0.69, respectively (Figure S.2b,d). However, kurtosis has a strong correlation only because kurtosis itself is strongly correlated with skewness (*R*^2^ = −0.67), which arises from the assumed relationships in equations 5–6.

We hypothesized that skewness would substantially impact species responses to climate warming, as climate warming is directional (i.e., a north-shifting fundamental niche), and skewness measures the direction of the heavy tail of the *R*_0_(*T*) curve. While kurtosis affects slope of the *R*_0_(*T*) curve near the tolerance limits, and this will affect the magnitude of the species’ response to climate warming, we do not expect to find qualitative effects of kurtosis on climate change impacts, as we have found for skewness.

**Figure S.2:**
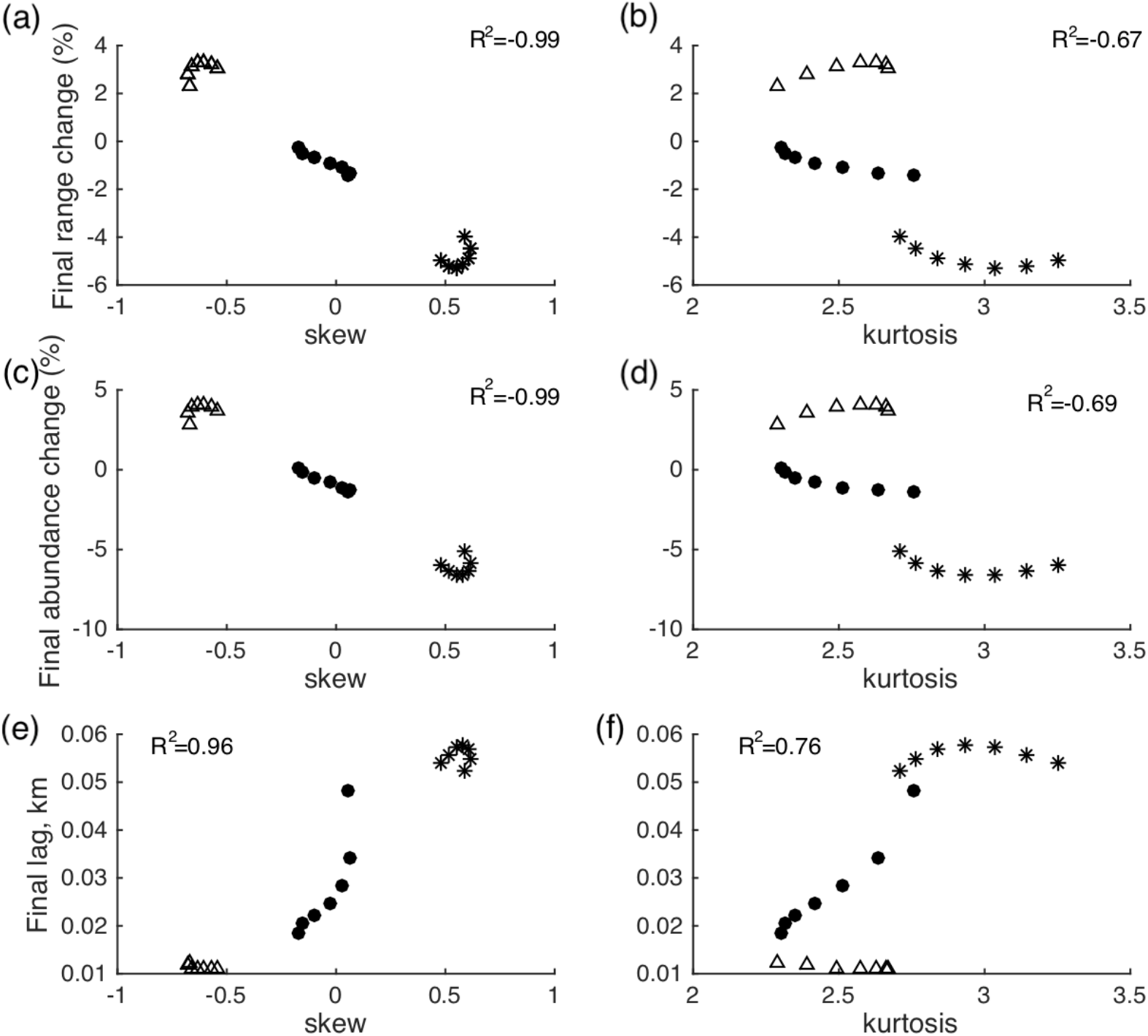
*R*_0_(*T*) skewness (left column) is strongly correlated with climate warming impacts, whereas *R*_0_(*T*) kurtosis (right column) is correlated with the impact of climate warming only since kurtosis is correlated with skewness (*R*^2^ = −0.67). The impacts of climate warming are measured as the final range change (top row), final abundance change (middle row), and final lag (bottom row). The symbols are cold-skew (*), warm-skew (Δ), and symmetric (•) fitness curves.

#### 3.2 Fitness curve rotations

To further test that it is indeed skewness (and not some other characteristic of *R*_0_(*T*)) that determines how range size, abundance, and lag change, we rotated *R*_0_(*T*) curves around a vertical axis to reverse skewness but maintain the shape otherwise, i.e., a rotated cold-skewed curve becomes a warm-skewed curve, and vice versa. Figure S.3 confirms that these rotations reverse results with respect to range size, abundance, and lag, in each case as expected.

**Figure S.3:**
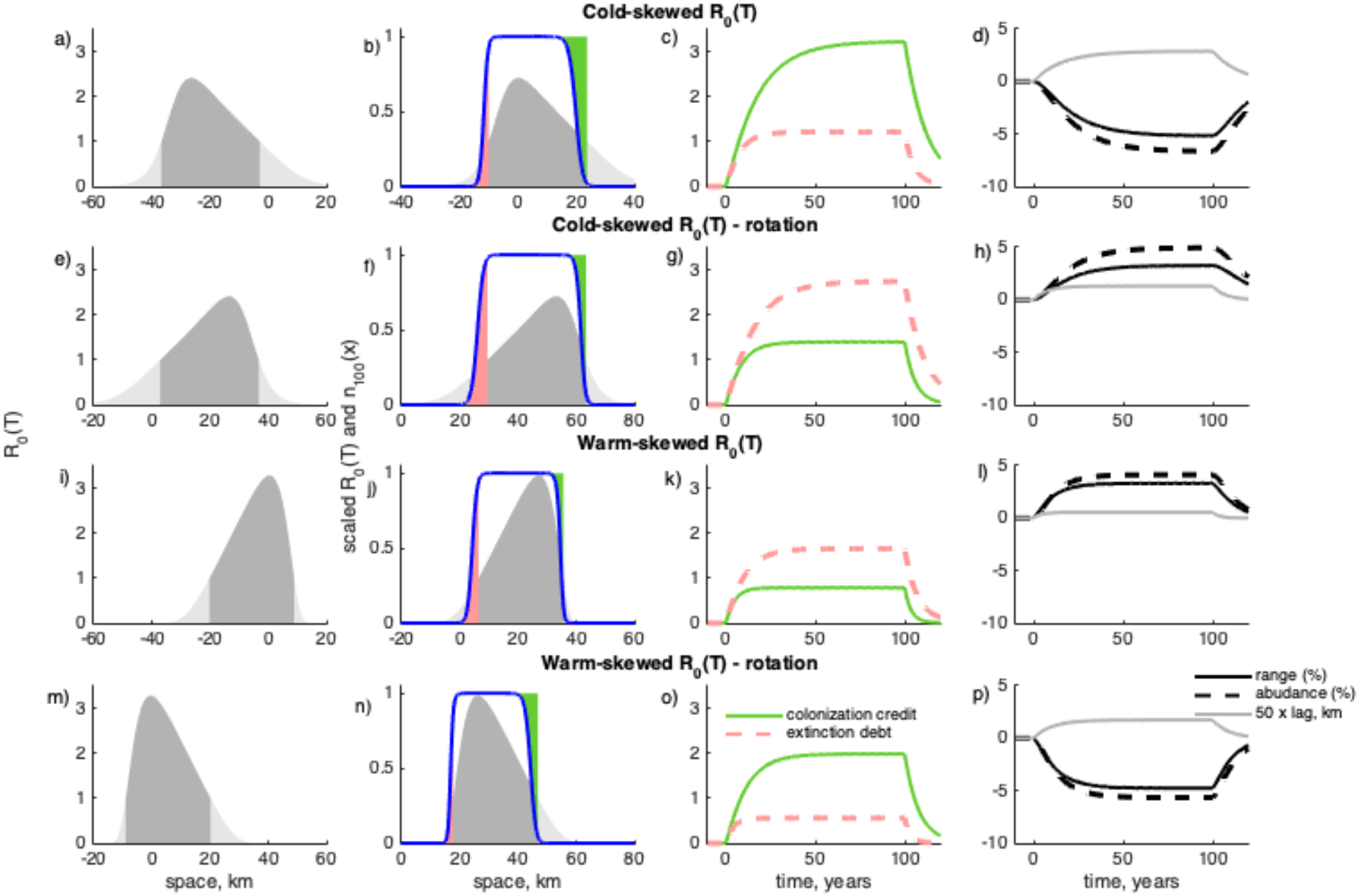
Fitness curves with the same skewness generate the same qualitative results whether they are generated by equations 4–7 in the main text or by rotation around a vertical axis. A cold-skewed fitness curve (as in Figure 3a) shown unchanged (a) and rotated around a vertical axis to result in a warm-skewed *R*_0_(*T*) (e). For a cold-skewed fitness curve rotated around a vertical axis, the climate warming impacts are qualitatively the same as those for a warm-skewed fitness curve: extinction debts exceed colonization credits (g), resulting in increased range size and abundance, and small lags (h). Similarly, for the warm-skewed fitness curve (i, as in Figure 3b) when rotated around a vertical axis to produce a cold-skewed fitness curve (m), the impacts of climate warming are qualitatively the same as for a cold-skewed fitness curve: colonization credits exceed extinction debts (o), resulting in decreased range size and abundance, and large lags (p). a),b),e),f),i),j),m),n) shows *R*_0_(T) > 1 in dark grey and *R*_0_(*T*) < 1 in light grey shading. b),f),j),n) shows for t= 100, the population density (blue curve), *R*_0_(*T*) (grey shaded region; multiplied by 0.3 for visual comparison), colonization credit (the area of the green shaded region), and extinction debt (the area of the pink shaded region).

#### 3.3 Climate cooling scenarios

We would expect our explanation of our climate warming results to also apply to a hypothetical climate cooling scenario. To test this, we assumed climate cooling of 0.1°C per year. As expected, all results are reversed relative to the warming scenario: populations with cold-skewed *R*_0_(*T*) are positively impacted by climate cooling, experiencing range size and abundance increases; by contrast, populations with warm-skewed *R*_0_(*T*) are negatively impacted, experiencing range size and abundance decreases (Figure S.4d,h). For climate cooling scenarios, the leading edge of the population density is to the south and the results shown in Figure S.4 confirm our reasoning that the shape of the fitness curve at the leading edge of the population density is key to understanding how a population will be affected by a shifting fundamental niche.

**Figure S.4:**
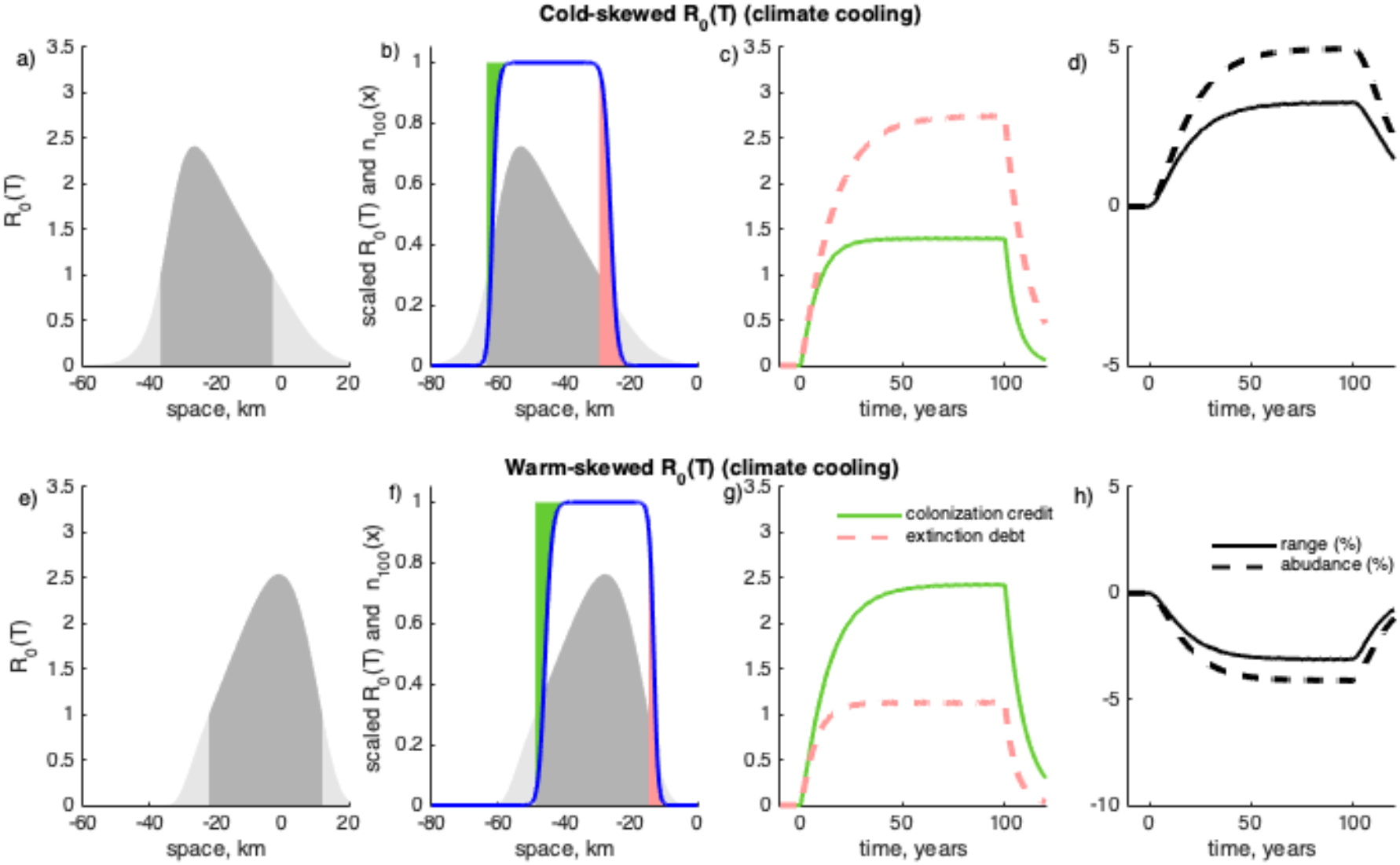
As expected, our results for climate warming scenarios are reversed for climate cooling scenarios: populations with cold-skewed *R*_0_(*T*) are positively impacted by climate cooling, while populations with warm-skewed *R*_0_(*T*) are negatively impacted. At *t* = 0, populations with cold- and warm-skewed fitness curves (as in Figure 3a,b in the main text) are subjected to climate cooling of 0.1°C per year. b) and d) show, at *t* = 100, the population density (blue curve), the fitness curve (multiplied by 0.3 for visual comparison; *R*_0_(*T*) > 1 dark grey and *R*_0_(*T*) < 1 light grey) and the corresponding colonization credit (the area of the green shaded region) and extinction debt (the area of the pink shaded region). Populations with cold-skewed fitness curves exhibit increased range size and abundance due to climate cooling (d), while populations with warm-skewed fitness curves experience decreased range and abundance (h).

### 4 Spatio-temporal dynamics at the trailing edge and extinction debts

In this section, we examine the spatio-temporal dynamics at the trailing edge of the population density to understand why extinction debts are generally slightly larger for populations with warm-skewed fitness curves. Figure S.5 compares the dynamics at the trailing edge by shifting the location of the warm tolerance limit for the population with the warm-skewed fitness curve by 16.55 kms to the south to align with the warm tolerance limit for the cold-skewed fitness curve (black dashed lines). By definition extinction debts occur to the south of the warm tolerance limit (Figure S.5, black dashed lines), and in this region both the population density (Figure S.5a) and the values of *R*_0_(*T*) (Figure S.5c) are larger for warm-skewed fitness curves (red) relative to cold-skewed fitness curves (blue), explaining why extinction debts are also larger (Figure S.5b). One reason why populations with cold-skewed fitness curves, in response to climate warming, have larger decreases in range size and abundance relative to warm-skewed fitness curves, is that while there are some regions in space where the values of *R*_0_(*T*) are larger for cold-skewed (Figure S.5d, blue) relative to warm-skewed fitness curves (Figure S.5d, red), for much of these regions the population density is either at carrying capacity or near zero, such that these larger *R*_0_(*T*) values do not translate into an increase, or a reduction in the decrease, of the range size.

**Figure S.5:**
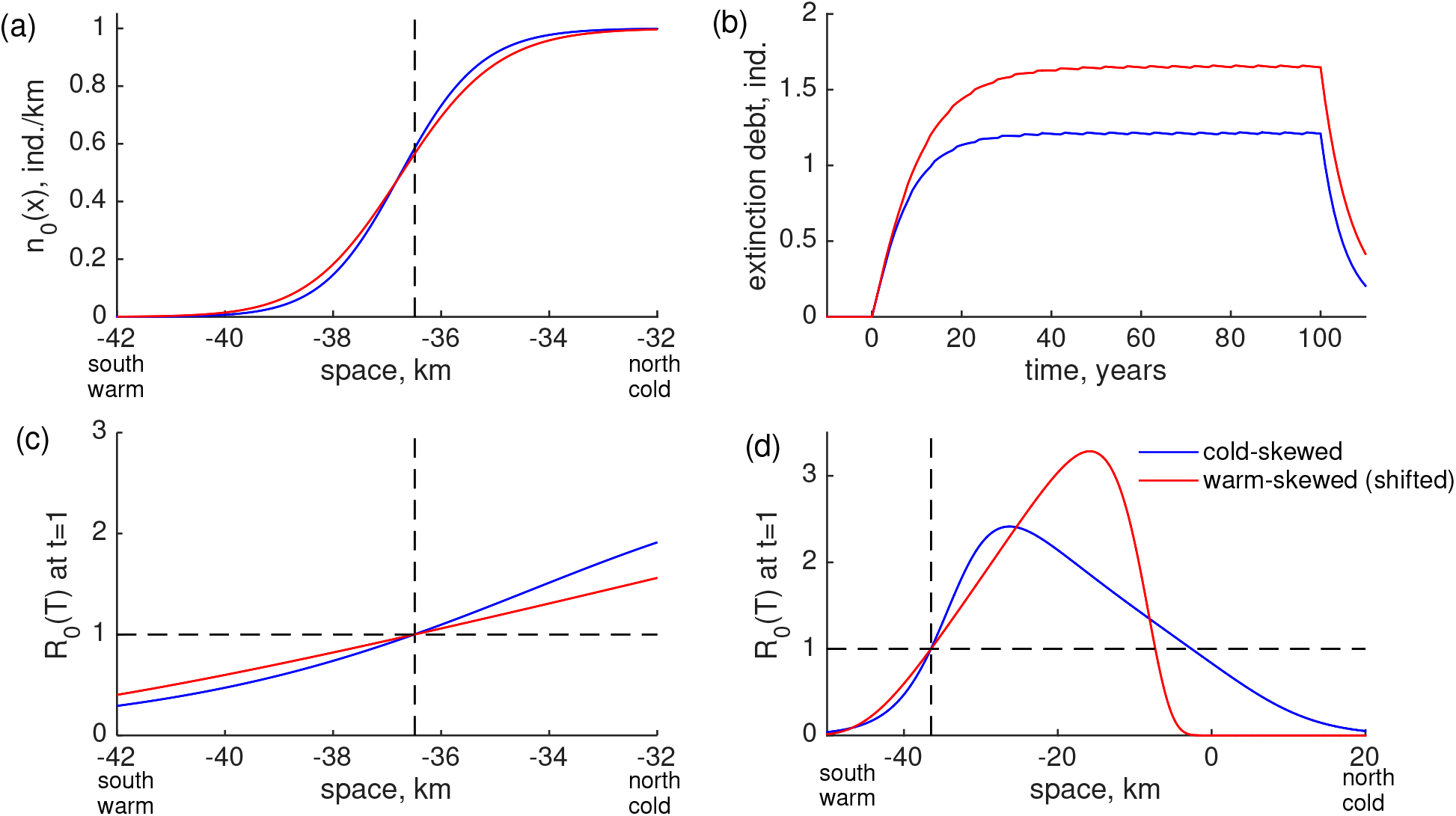
Extinction debts are larger for warm-skewed fitness curves relative to cold-skewed fitness curves because the population density and fitness curve values are larger for warm-skewed fitness curves for all locations south of the warm tolerance limit. The extinction debt at t =1 is defined as the numerical integral of the population density over all locations south of the warm tolerance limit, which, for *t* = 1, is marked with the dashed lines (a,c,d). The population density south of the warm tolerance limit at *t* = 1 depends on the population density at *t* =0 (a) and the values of *R*_0_(T) at *t* = 1 (c). At all points south of the warm tolerance limit, the population density at *t* = 0 (a) and values of *R*_0_(*T*) at *t* = 1 (c) are greater for populations with warm-skewed fitness curves (red) relative to cold-skewed fitness curves (blue), resulting in a larger extinction debt, at *t* = 1, for populations with warm-skewed fitness curves. This pattern of larger extinction debts for warm-skewed fitness curves, occurs not only for *t* = 1, but for all times when climate warming occurs (b, i.e., from *t* = 0 to 100). As for c), d) shows *R*_0_(*T*) at *t* = 1, but d) shows a larger region of space to show the complete shape of the fitness curves. To facilitate comparisons with the cold-skewed fitness curve, the warm-skewed fitness curve and population density shown in a),c), and d) are shifted 16.55km to the south so that the location of the warm tolerance limit aligns with the warm tolerance limit of the cold-skewed fitness curve. All parameters are as for Figure 3a,b in the main text.

### 5 Effect of dispersal distance

In this section, we analyze how mean dispersal distance affects a population’s colonization credits, extinction debts, range size, abundance, and lag during range changes.

#### 5.1 Colonization credit and extinction debt

Colonization credit occurs in locations where the population is below its carrying capacity despite *R*_0_(*T*) > 1, and is an abundance quantified as the difference between the carrying capacity and the population density numerically integrated across locations meeting these criteria. Colonization credit quantifies the additional abundance the environment could support, if it were not for dispersal limitation. In contrast, extinction debt occurs in areas where the population remains temporarily present despite *R*_0_(*T*) < 1, and is defined as the numerical integral of the population density across all these locations. Extinction debt is thus the abundance of this population that can only persist temporarily (see Figure 1 in the main text).

Figure S.6 shows that both colonization credits and extinction debts decrease with mean dispersal distance. The decrease in colonization credit as the mean dispersal distance increases (Figure S.6, green solid lines) is due to an increasing proportion of offspring dispersing longer distances from their natal sites, and are thus better able to colonize new habitat and keep pace with their moving fundamental niche. There are two factors that explain why extinction debt decreases with increasing mean dispersal distance (Figure S.6, pink dashed lines). Firstly, the population is better able to track the location of the fundamental niche with larger dispersal distances, meaning that fewer individuals begin their dispersal from a position behind the trailing edge of the niche, resulting in smaller extinction debts. Secondly, any group of individuals that does start behind the trailing edge, with larger dispersal distances, will see a larger proportion of dispersers catching up with the niche, again lessening the extinction debt. Despite both colonization credit and extinction debt decreasing with mean dispersal distance, populations with cold-skewed fitness curves remain more adversely affected by climate warming (Figure S.6a) in comparison to their warm-skewed counterparts (Figure S.6b) for all choices of mean dispersal distance (Figure S.6). Figure S.6 shows that if colonization credit is greater than extinction debt, then this holds for all values of the mean dispersal distance (and visa versa), which in turn implies that abundance changes for fitness curves of a given skewness are robust to different mean dispersal distances.

**Figure S.6:**
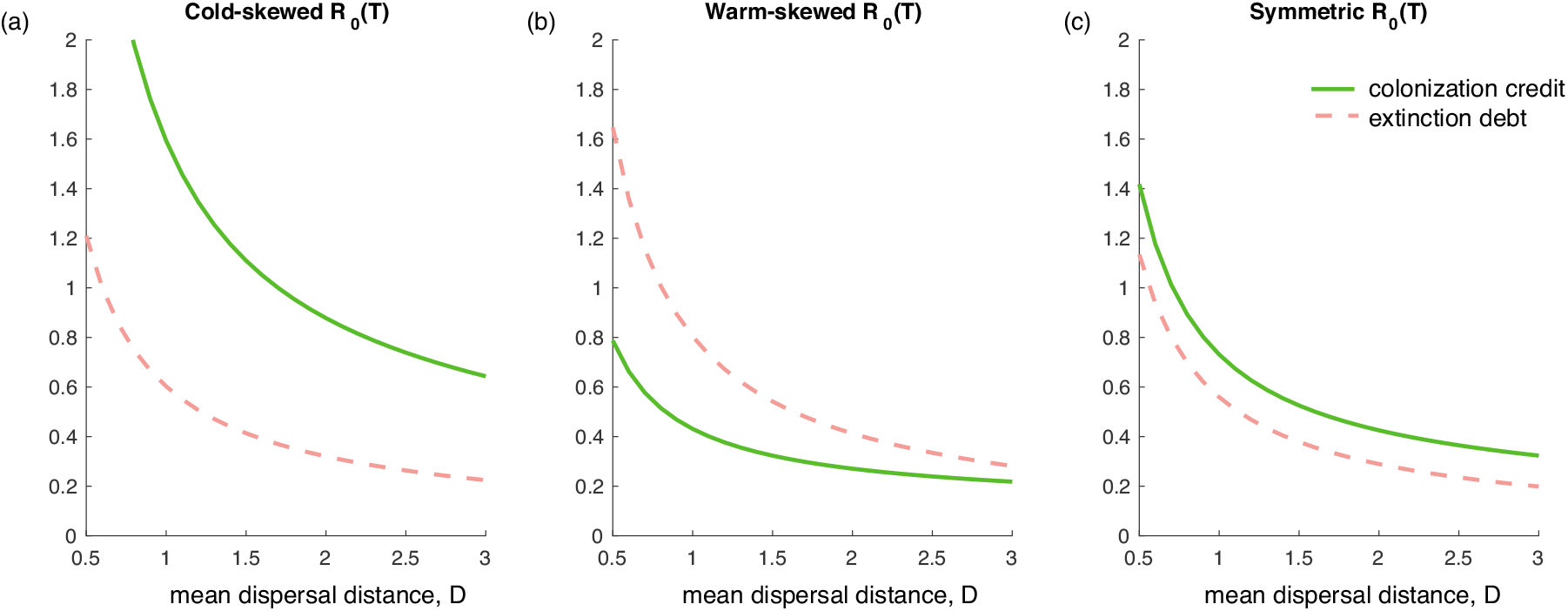
As the mean dispersal distance increases, colonization credits and extinction debts (after 100 years of climate warming) decrease. Simulation parameters are the same as for Figure 4 of the main text, except that the mean dispersal distance is varied as indicated on the *x*-axis.

#### 5.2 Final range size, final abundance, and final lag

Populations are better able to keep pace with climate change as mean dispersal distance (*D*) increases, resulting in smaller lags, and larger final range sizes and final abundances (Figure S.7). Our qualitative results from the main text remain unchanged for different choices of mean dispersal distance (Figure S.7): climate warming results in range size and abundance decreases for organisms with cold-skewed fitness curves and in range size and abundance increases for organisms with warm-skewed fitness curves (Figure S.7). Mean dispersal distances smaller than 0.5 km were not considered in our simulations as the population with the cold-skewed fitness curve fails to keep pace with climate change in these cases.

**Figure S.7:**
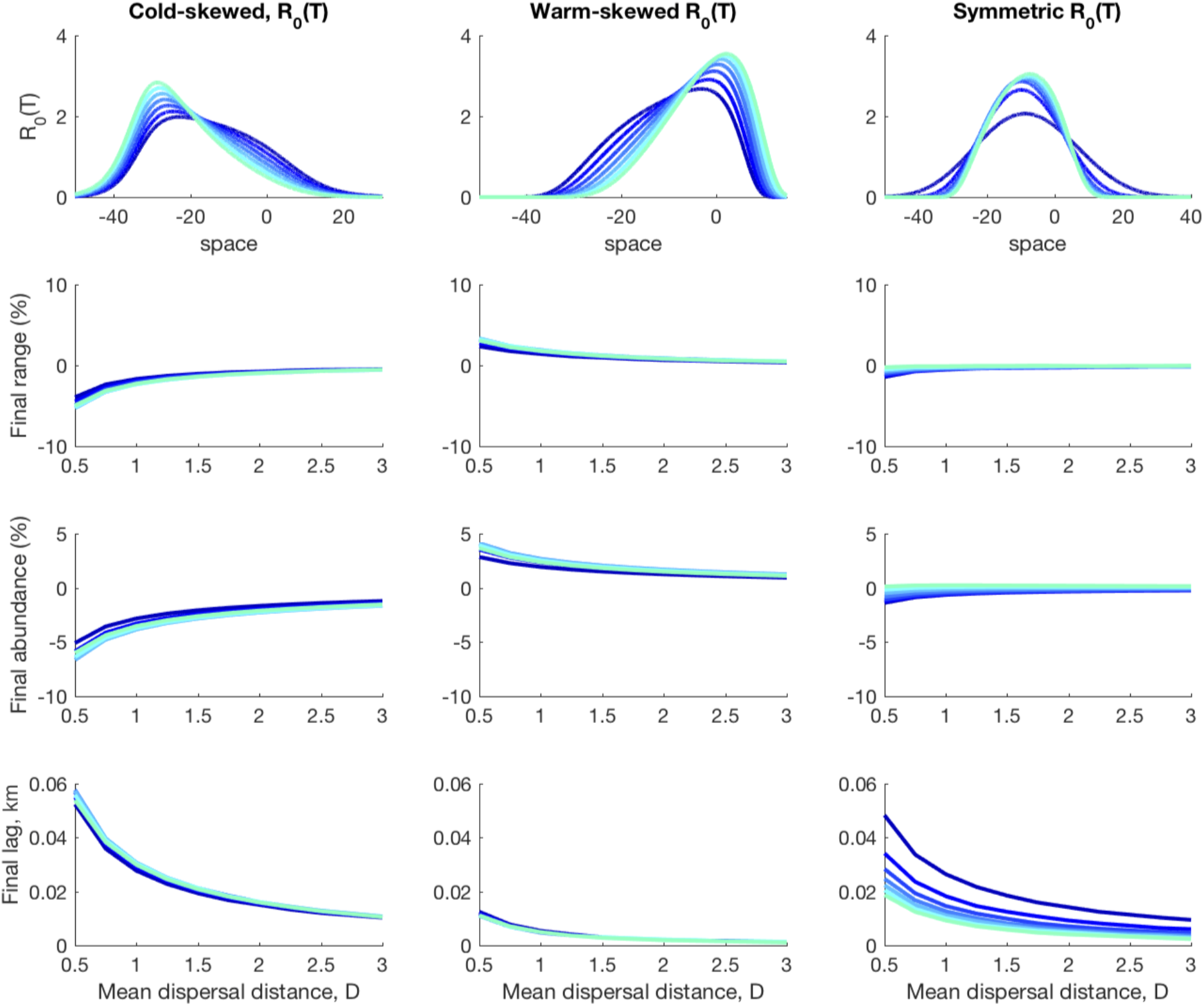
The impacts of climate warming are qualitatively insensitive to changes in the mean dispersal distance in the Laplace dispersal kernel: a larger mean dispersal distance lessens the impacts of warming, but the direction of change is maintained for all cases of *R*_0_(*T*) skewness. Parameters are identical as in Figure 4 of the main text, but in this figure only the final range size, final abundance and final lag (at *t* = 100) are shown.

#### 5.3 Asymmetric dispersal

In the main text (equation 2), we assume that offspring are equally likely to disperse to the north and to the south. However, species dispersal may be biased in a particular direction. To test the sensitivity of our results to our assumption that dispersal is symmetric, we consider an assymetric dispersal kernel,

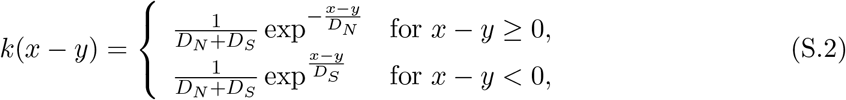

where *D_N_* >*D_S_* gives north-biased and *D_S_* >*D_N_* gives south-biased dispersal. The probability of dispersing to the north is *DN* /(*DN* + *DS*), and the mean dispersal distance when dispersal is to the north is 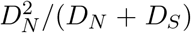, with analogous formulas for the probability of southward dispersal and the mean dispersal distance to the south.

For north-biased dispersal, where dispersal is biased in the direction of the shifting fundamental niche, we find that the negative impacts of climate warming are reduced and the positive impacts are increased (Figure S.8e-j; the mint curve lies above the cyan curve). For south-biased dispersal, where dispersal is biased away from the direction of the shifting fundamental niche, we find the opposite: the negative impacts of climate warming are increased and the positive impacts are decreased (Figure S.8e-j; the blue curve lies below the cyan curve).

Figure S.8 shows no qualitative changes in our main results regarding cold-skewed fundamental niches being negatively impacted by climate warming and warm-skewed fundamental niches being positively impacted (summarized in Table 1 of the main text). It was not possible to consider a stronger southward dispersal bias because in doing so, the population was unable to keep pace with climate warming.

**Figure S.8:**
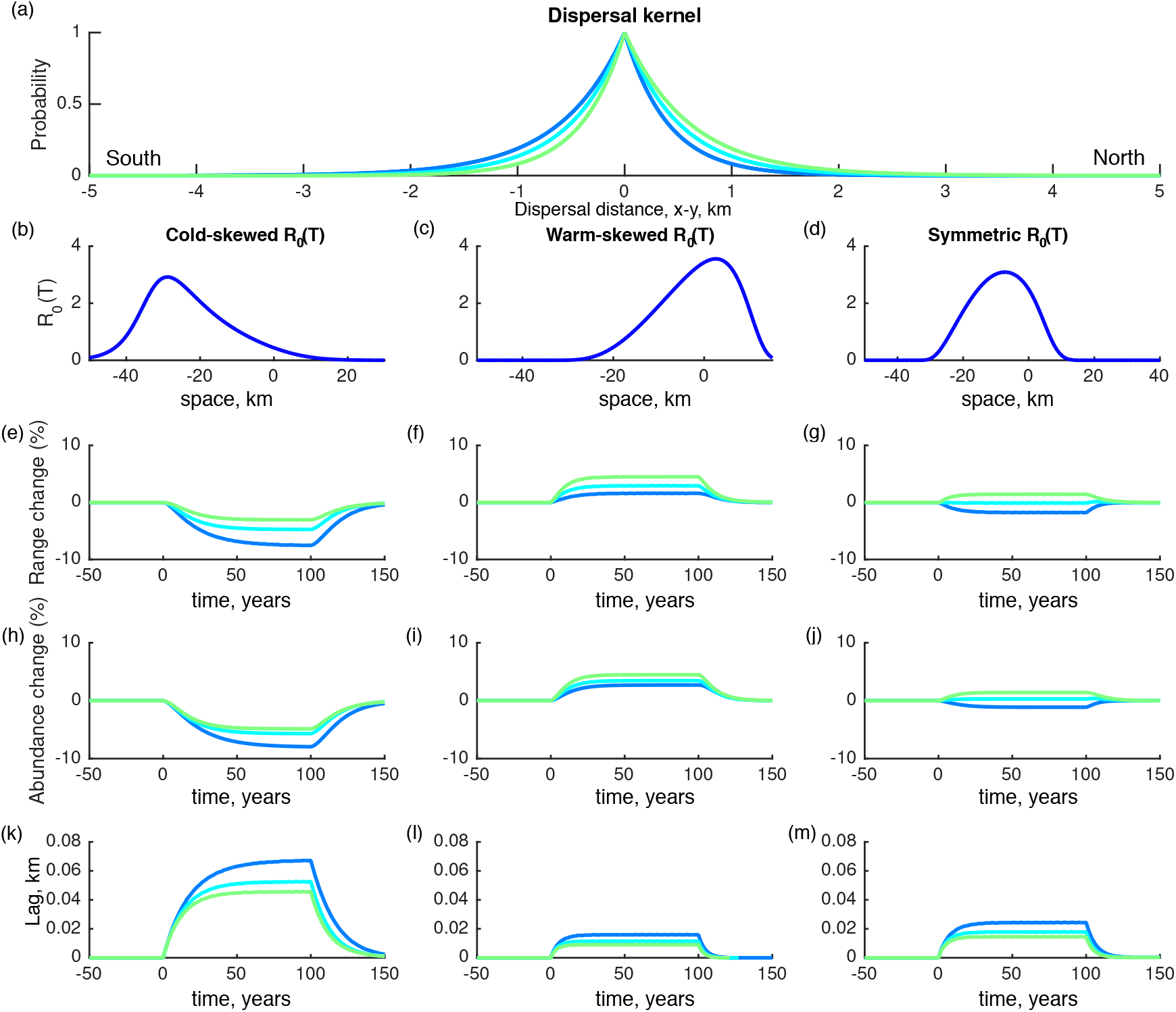
For asymmetric dispersal kernels (equation S.2) our conclusions that populations with cold-skewed *R*_0_(*T*) experience range and abundance losses, while populations with warm-skewed *R*_0_ (*T*) experience gains is unchanged. (a) The dispersal kernels that we consider are Laplace-type distributions (i.e double exponential distributions), but the mean dispersal distance to the north is not necessarily equal to that of the south. The mint green curve shows north-biased dispersal because *D_N_* =0.8 > 0.2km=*D_S_*. The cyan curve shows non-biased dispersal with *D_N_* = *D_S_* =0.5 km. The blue curve has south-biased dispersal, with *D_N_* =0.2 < 0.8 km = *D_S_*. (b-d) Rather than the full range of *R*_0_(*T*) skewnesses that are considered in the main text (Figure 4), here we consider only the most cold- and warm-skewed curves by setting *E_ρ_* = 1.2 and *E_μ_* = 0 eV (b), *E_ρ_* = 0 and *E_μ_* = 1.2 eV (c), and *E_ρ_* = *E_μ_* = 1.2 eV (d). Populations with cold-skewed *R*_0_(*T*) experience range (e) and abundance decreases (h) while populations with warm-skewed *R*_0_(*T*) experience increases (f,i) consistent with the findings we reported in Table 1 of the main text.

### 6 Simulations using the Ricker growth function

In the main text, we evaluated range change dynamics for the case of compensatory density dependence using a Beverton-Holt population growth model (equation 3). Here, we assess whether results differ for overcompensatory density dependence using the Ricker model for population growth. The Ricker growth function is given by,

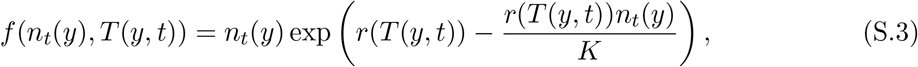

where *r*(*T*(*t,y*)) = ln(*R*_0_(*T*(*y,t*))) and *K* is the carrying capacity. As in the Beverton-Holt scenario, this carrying capacity is independent of *R*_0_, as can be seen from rewriting equation S.3 as,

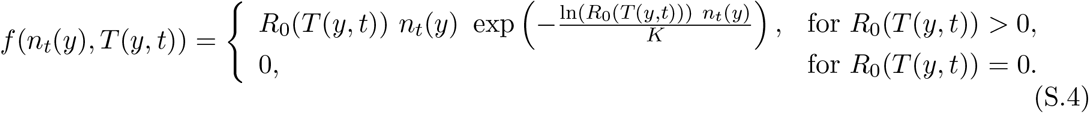

Combining the Ricker growth function with the Laplace dispersal kernel yields near identical values for our summary statistics as in the Beverton-Holt/Laplace scenario (Figures S.9 and S.10). The only exception is for large values of *R*_0_(*T*) as discussed in the next section.

**Figure S.9:**
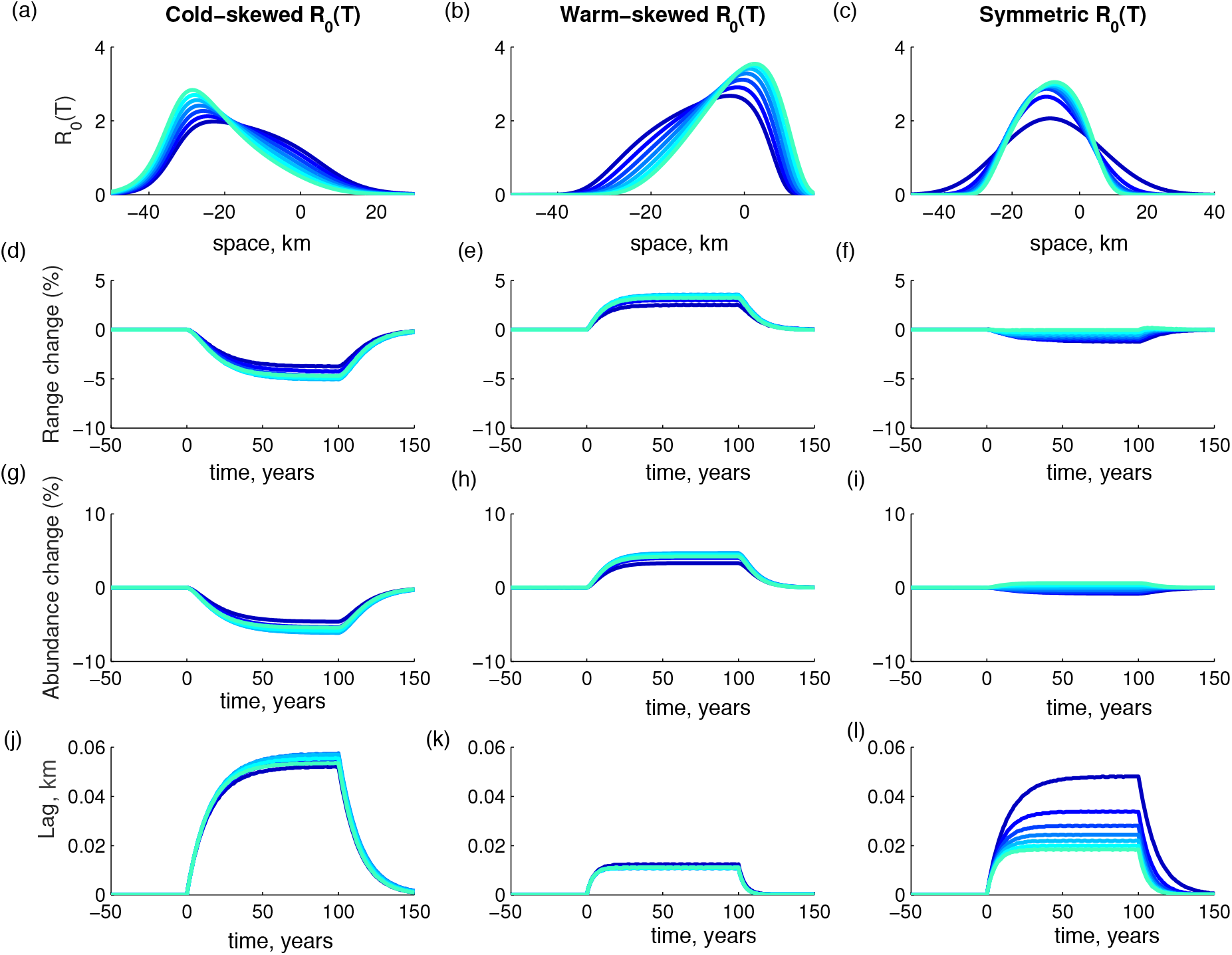
Analogous to Figure 4 in the main text, but now using the Ricker population growth model instead of the Beverton-Holt model.

**Figure S.10:**
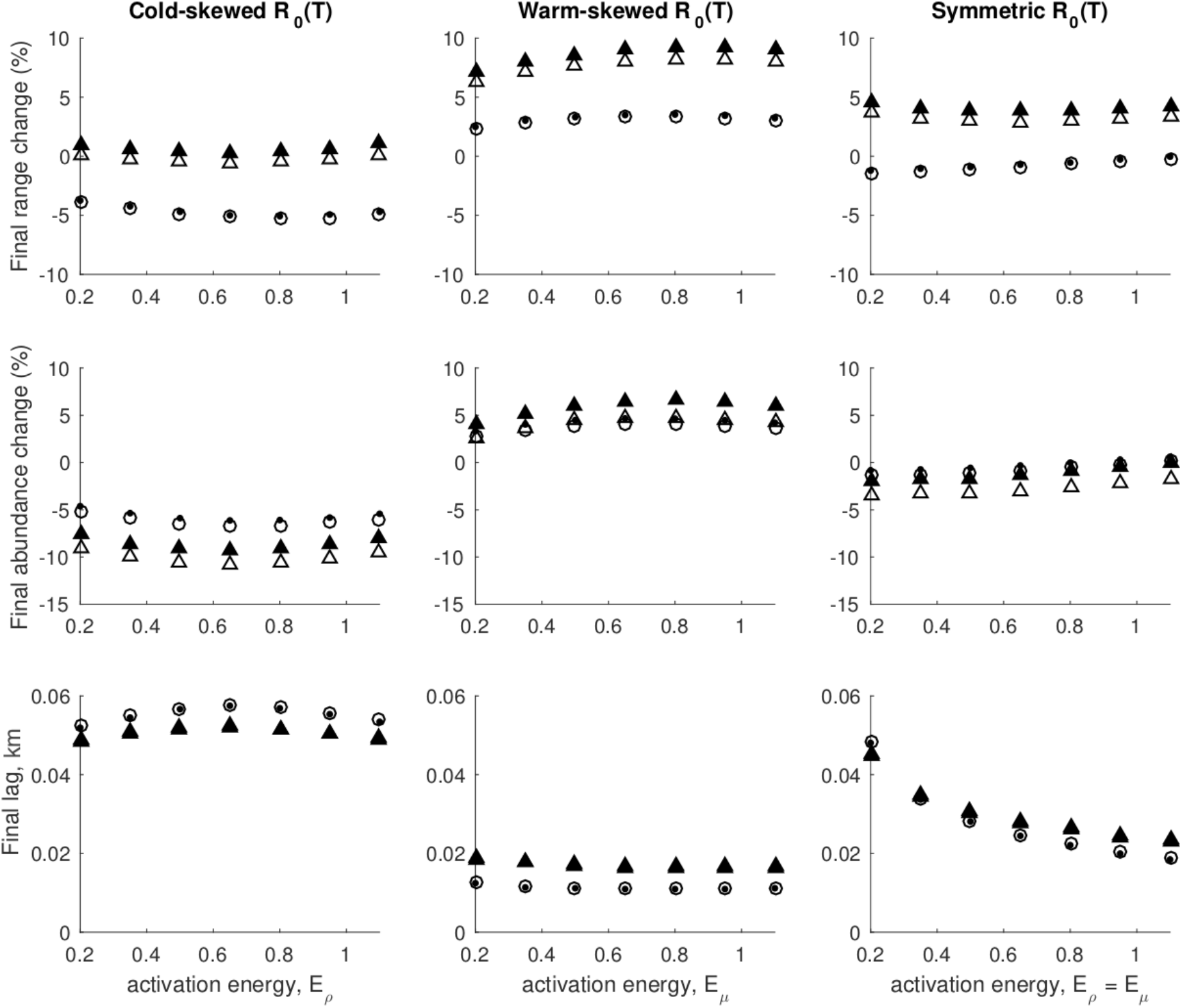
The effect of *R*_0_(*T*) skewness on a population’s response to climate warming is similar for different combinations of Beverton-Holt or Ricker population growth models and Laplace or Cauchy dispersal kernels. The final range size (top row), final abundance (middle row) and final lag (bottom row) after 100 years of climate warming are shown for the Beverton-Holt/Laplace (o), Ricker/Laplace (·), Beverton-Holt/Cauchy (Δ), and Ricker/Cauchy (▲) combinations of population growth models and dispersal kernels. In each of these scenarios, populations with cold-skewed *R*_0_(*T*) are adversely impacted by climate warming, warm-skewed *R*_0_(*T*) benefit from climate warming, and symmetric *R*_0_(*T*) are unaffected or experience mixed responses. Simulation parameters are as in Figure 4 in the main text and *β* =0.005 km was used for the Cauchy dispersal kernel. In the bottom row, the Beverton-Holt/Cauchy (Δ) and the Ricker/Cauchy (▲) scenarios are visually indistinguishable.

#### 6.1 Population cycles

Climate change may dampen or eliminate population cycles. For large values of *R*_0_(*T*), prior to climate change the Ricker population growth function gives rise to a two-cycle in population density (Figure S.11a), but when climate warming occurs the population cycles dampen (Figure S.11b,d). After the end of climate warming, the population density returns to an identical two-cycle pattern, but displaced northwards due to the warming-induced northern shift in the fundamental niche.

**Figure S.11:**
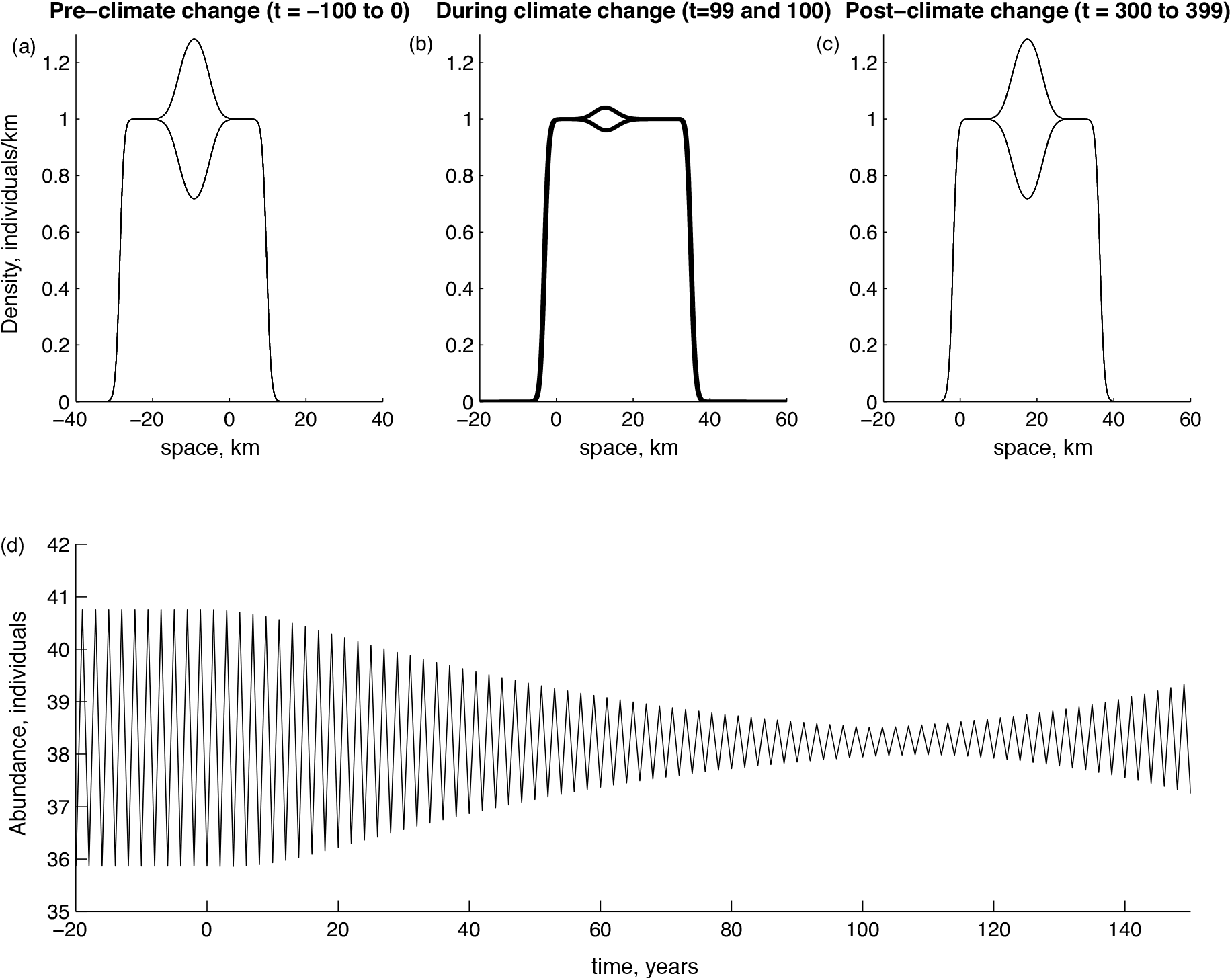
Climate warming may dampen population cycles arising from overcompensation. (a) For Ricker population growth, prior to climate warming the population exhibits a stable two-cycle (the population density is shown every year for 100 years prior to the onset of climate warming from t=-100 to 0). (b) The population cycles are dampened during climate warming and the abundance becomes less variable (d). (c) After climate warming ends the population converges back to the original stable two-cycle, but displaced northwards. The parameters are *D* = 0.5 km, 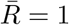, *ρ*_0_=44 with all other parameters as in Figure 4 of the main text.

### 7 Simulations using the Cauchy dispersal kernel

In the main text, we evaluated range change dynamics for the case of an exponentially bounded (Laplace) dispersal kernel. Here, we assess whether results differ for a fat-tailed dispersal kernel (Cauchy) because population spread rates are known to be greatly influenced by the tail of the kernel (Zhou et al., 2013). The Cauchy kernel is commonly used to model ‘fat-tailed’ dispersal and is given by,

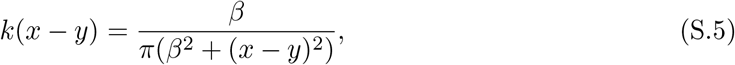

where *β* is a shape parameter. Compared to the Laplace distribution, which exhibits exponential decay in the tail of the distribution, the Cauchy distribution is ‘fat-tailed’ due to the power law decay of the tail. The associated higher number of long-distance dispersal events explains both why range sizes are larger under a Cauchy kernel and why abundances are smaller (more individuals are lost due to long-distance dispersal into hostile habitat; Figure S.10). Nevertheless, with the Cauchy dispersal kernel our general result remains: climate warming will adversely affect populations with cold-skewed fitness curves (substantially decreasing abundance), benefit populations with warm-skewed fitness curves, and will have mixed effects on populations with symmetric fitness curves.

Simulations that use the Cauchy dispersal kernel were run with *β* =0.005 km, where 2*β* is the width of the distribution at half its maximum (for reference, the corresponding width for the Laplace distribution is 2*D* ln 2, where *D* is the mean dispersal distance). Figure S.12 is analogous to Figure 4 of the main text except that the Cauchy dispersal kernel is used. Ricker population growth combined with a Cauchy dispersal kernel yields near identical results to the combination of the Beverton-Holt growth function with the Cauchy dispersal kernel (Figure S.10, triangle symbols).

**Figure S.12:**
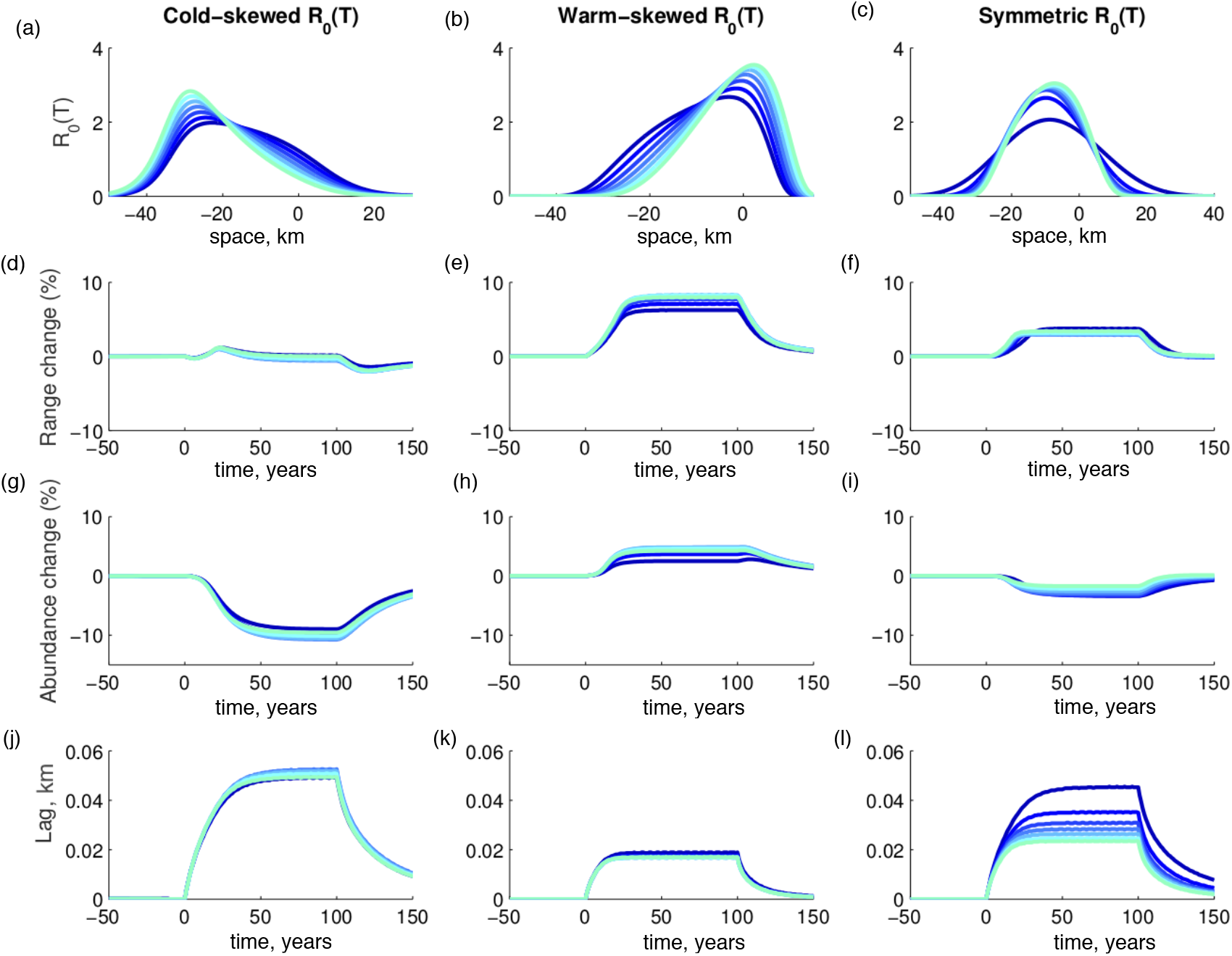
This figure is analogous to Figure 4 in the main text except that the Cauchy dispersal kernel replaces the Laplace dispersal kernel from Figure 4. In contrast to the results found for the Laplace dispersal kernel, for the Cauchy dispersal kernel we observe: 1) larger range sizes, and 2) smaller abundances during climate change. For this figure *β* =0.005 km and all other parameters are identical to Figure 4.

### 8 Stochastic fluctuations in annual temperature

In the main text, we analyzed the impacts of climate warming, assuming a unidirectional temperature increase of 0.1°C per year at all locations. Here, we relax this simplifying assumption and evaluate how annual stochastic variation around the unidirectionally increasing mean temperature affects our results. We implemented stochastic changes in climate warming by adding a uniformly distributed random variable to the annual temperature change. Specifically, we set the temperature gradient at time, *t*, as,

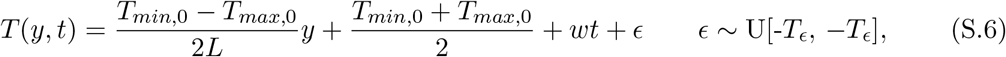

where [─*T_ϵ_, T_ϵ_*] is the range of the uniform distribution and all other parameters are defined as in Section 1, where the temperature gradient is originally discussed. The stochastic effects of warming occur each year and apply equally to all points in the spatial temperature gradient. Historical temperature data suggests that the annual difference between predicted and observed temperature is no more than ±0.3°C (NASA’s Goddard Institute for Space Studies, 2019). We consider moderate (*T_ϵ_* = 1°C) and extreme (*T_ϵ_* = 2.5°C) fluctuations, but we note that even our moderate fluctuation simulations (*T_ϵ_* = 1°C) are more variable than the observed annual temperature variability of less than ±0.3°C.

We find that our deterministic model approximates the central tendency of the impacts of climate warming for our simulations with moderate stochasticity (Figure S.13, red and blue lines). For extreme temperature fluctuations, all populations are negatively affected, regardless of their fitness curve skew (Figure S.14e-j, red lines; from *t* = 0 to 100 the range size and abundance decreases), because large temperature shifts imply large shifts in the locations of suitable habitats, thereby leaving part or all of a population within unsuitable habitat intermittently.

For our simulations with moderate stochasticity (Figure S.13) our qualitative conclusions are maintained, except that populations with symmetric fitness curves are negatively impacted by climate warming, where they might otherwise have been unaffected (Table 1, in the main text). For the warm-skewed fitness curve, we see the beneficial effects of climate warming still remain (Figure S.13 f,i; increases in range size and abundance from *t* = 0 to 100), but that stochasticity (red line) decreases the magnitude of this effect. For populations with cold-skewed fitness curves, considering a moderate level of stochasticity does not have an effect (Figure S.13 e,h,k; red lines: the mean of the stochastic simulations, and blue lines: the deterministic simulation, are nearly indistinguishable).

**Figure S.13:**
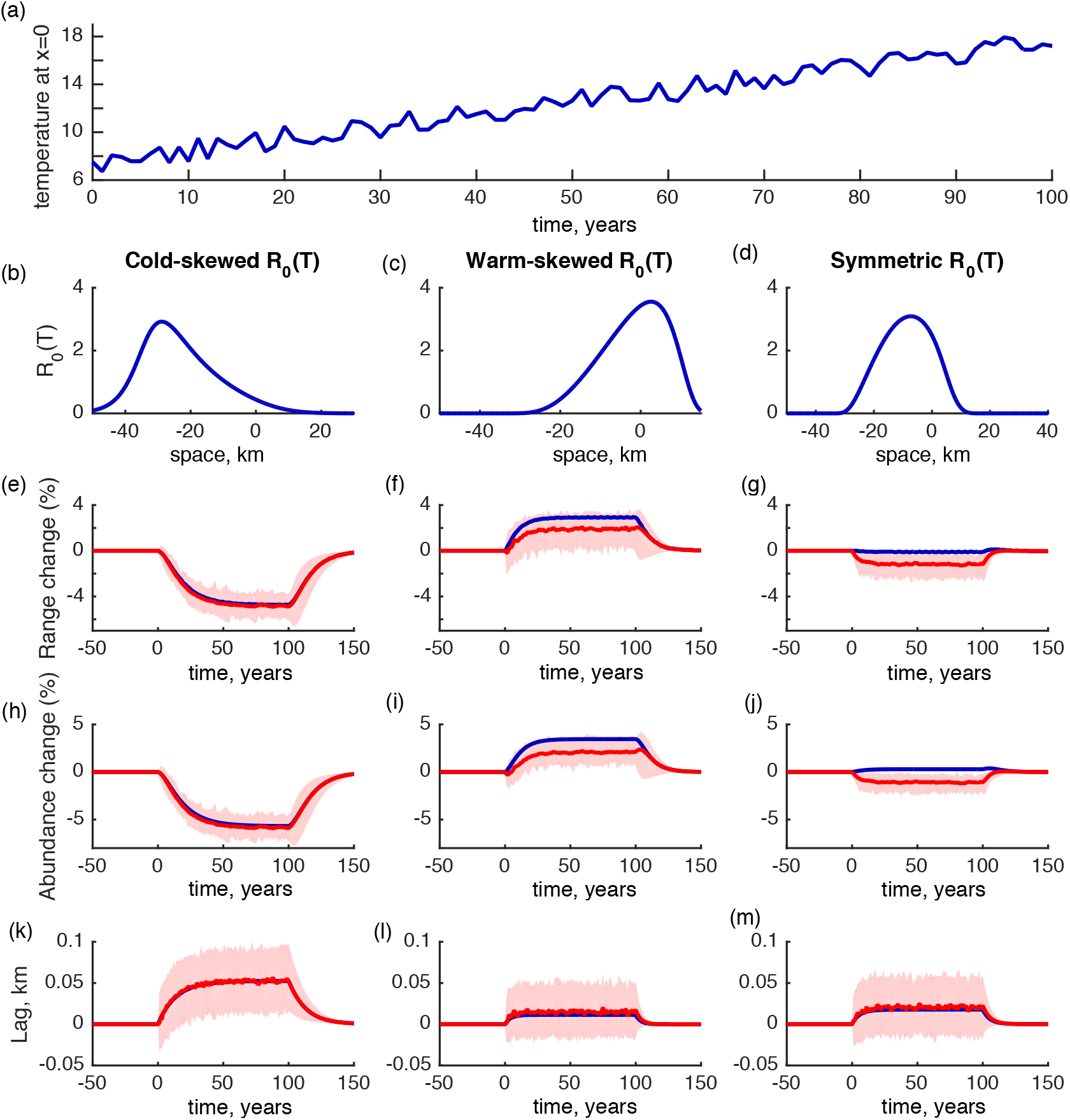
When annual stochastic variation around the unidirectionally increasing mean temperature is *T_ϵ_* = 1°C (see equation S.6), populations with warm-skewed fitness curves benefit from climate warming, while populations with cold-skewed fitness curves are adversely affected. (a) Temperature in degrees Celsius at *x* = 0 for one stochastic realization of climate warming. The cold-skewed (b), warm-skewed (c), and symmetric (d) fitness curves used for the results shown in (e-m), where (e-m) show the mean (red line) and range (red shaded) for 100 realizations of how range, abundance, and lag change when climate is stochastic, as compared to when climate is assumed to be deterministic (blue). Stochasticity does not appreciably change our results for populations with cold-skewed fitness curves (i.e. the blue and red lines are indistinguishable in e,h and k). When stochasticity is included, populations with warm-skewed fitness curves still benefit as range size (f) and abundance (i) increase, however, this effect is less pronounced than when climate warming is deterministic (blue). Populations with symmetric fitness curves (g,j,m) are unaffected by climate warming when climate is deterministic (blue line), but are adversely affected when climate warming is stochastic (red line).

**Figure S.14:**
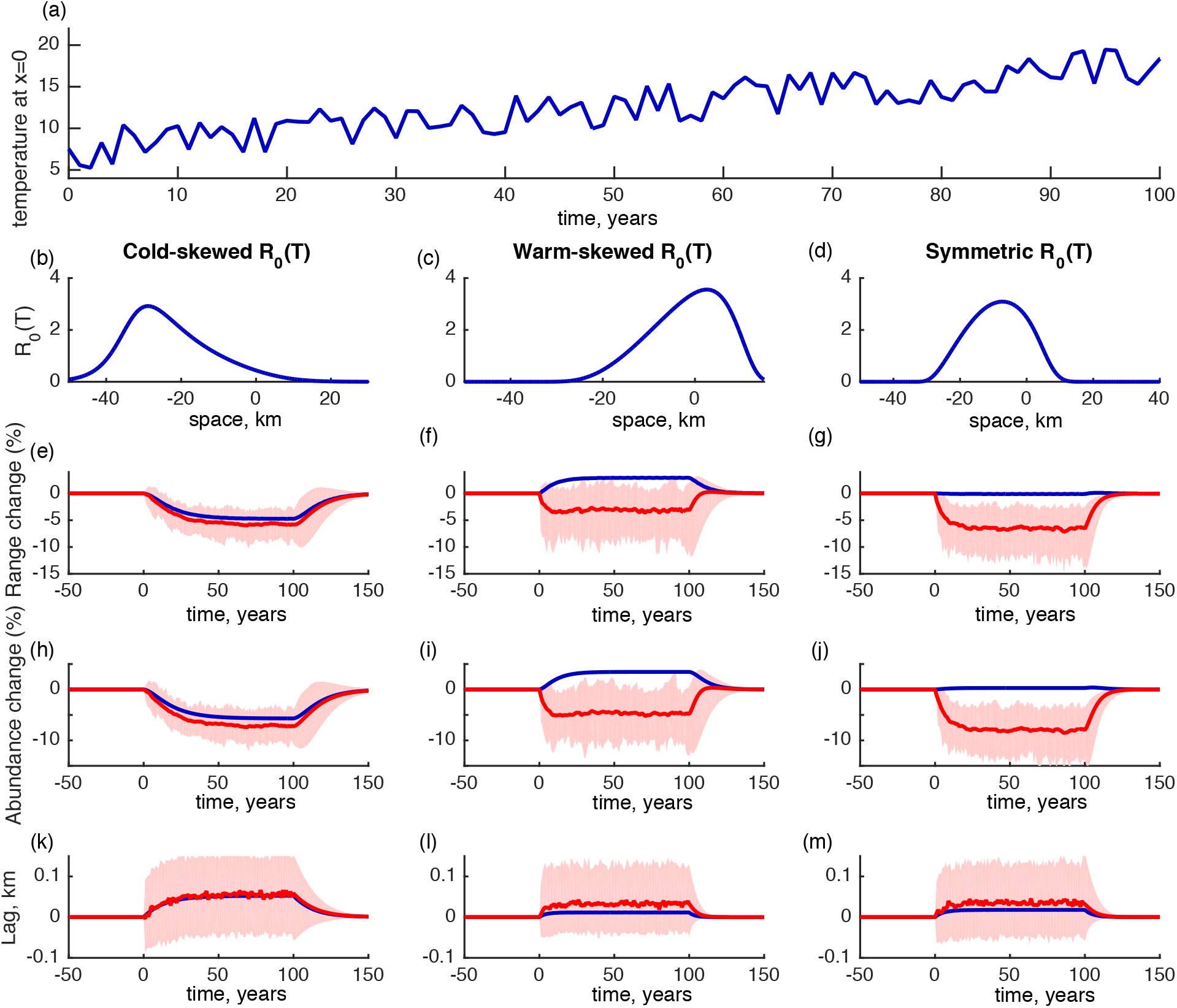
When annual stochastic variation around the unidirectionally increasing mean temperature is *T_ϵ_* = 2.5°C (see equation S.6), all populations are negatively impacted by climate warming. (a) Temperature in degrees Celsius at *x* = 0 for one stochastic realization of climate warming. The cold-skewed (b), warm-skewed (c), and symmetric (d) fitness curves used for the results shown in (e-m), where (e-m) show the mean (red line) and range (red shaded) for 100 realizations when climate is stochastic, as compared to when climate is deterministic (blue). For all the stochastic simulations (red lines in (e-m)) after the onset of climate warming at *t* =0 range size (e-g) and abundance (h-j) decreas, suggesting an adverse impact of climate warming. The qualitative effect of climate warming is reversed for warm-skewed fitness curves (f,i) under stochastic (red line) as compared to deterministic climate (blue line).

## Notes

#### Summary of Updates

The manuscript was reviewed at Proceedings of the Royal Society London B and revised to address the reviewers suggestions. The main text has had minor edits and additional analyses have been added to the Electronic Supplementary Material.

https://doi.org/10.6084/m9.figshare.6955370.v4

